# PDIP38 is a novel adaptor-like modulator of the mitochondrial AAA+ protease CLPXP

**DOI:** 10.1101/2020.05.19.105320

**Authors:** Philip R. Strack, Erica J. Brodie, Hanmiao Zhan, Verena J. Schuenemann, Liz J. Valente, Tamanna Saiyed, Kornelius Zeth, Kaye N. Truscott, David A. Dougan

## Abstract

Polymerase δ interacting protein of 38 kDa (PDIP38) was originally identified in a yeast two hybrid screen as an interacting protein of DNA polymerase delta, more than a decade ago. Since this time several subcellular locations have been reported and hence its function remains controversial. Our current understanding of PDIP38 function has also been hampered by a lack of detailed biochemical or structural analysis of this protein. Here we show, that human PDIP38 is directed to the mitochondrion, where it resides in the matrix compartment, together with its partner protein CLPX. PDIP38 is a bifunctional protein, composed of two conserved domains separated by an α-helical hinge region (or middle domain). The N-terminal (YccV-like) domain of PDIP38 forms an SH3-like β-barrel, which interacts specifically with CLPX, via the adaptor docking loop within the N-terminal Zinc binding domain (ZBD) of CLPX. In contrast, the C-terminal (DUF525) domain forms an Immunoglobin-like β-sandwich fold, which contains a highly conserved hydrophobic groove. Based on the physicochemical properties of this groove, we propose that PDIP38 is required for the recognition (and delivery to CLPXP) of proteins bearing specific hydrophobic degrons, potentially located at the termini of the target protein. Significantly, interaction with PDIP38 stabilizes the steady state levels of CLPX *in vivo*. Consistent with these data, PDIP38 inhibits the LONM-mediated turnover of CLPX *in vitro.* Collectively, our findings shed new light on the mechanistic and functional significance of PDIP38, indicating that in contrast to its initial identification as a nuclear protein, PIDP38 is a *bona fide* mitochondrial adaptor protein for the CLPXP protease.

## Introduction

Polymerase δ interacting protein of 38 kDa (PDIP38, also known as POLDIP2 and mitogenin) was originally discovered through a yeast two hybrid screen, as a p50 (subunit of DNA polymerase delta) interacting protein ^1^. Subsequently, PDIP38 has been identified in the nucleus, where it is proposed to play a role in DNA repair ^2-5^. It has also been located in the cytoplasm and the plasma membrane 2,4-6 where it has been implicated in a variety of cellular functions, ranging from cell proliferation ^7^, to regulation of the extracellular matrix ^8^, oxidative signalling and cell migration^9^, Tau aggregation^10^ and cancer^11^. Despite the extensive repertoire of putative physiological functions for PDIP38, few of the proposed interactions have been biochemically validated. Indeed, in contrast to its proposed role in the nucleus, there is growing evidence to suggest that PDIP38, through its interaction with CLPX, forms part of the mitochondrial protein homeostasis (proteostasis) network ^12-14^. However, currently little is known about the structure or function of PDIP38, its mechanism of action or its role in mitochondrial proteostasis.

Proteostasis involves the constant surveillance and maintenance of the proteome, from a proteins synthesis on a ribosome, through its folding and transport to the correct subcellular location and ultimately its removal from the cell in a timely manner^15^. This process is maintained by a network of proteins, which includes proteolytic machines and their associated cofactors that are responsible for the timely recognition and removal of damaged or unwanted proteins. In eukaryotes, the major non-lysosomal degradation pathway is mediated by the Ubiquitin (Ub) Proteasome System (UPS), which is responsible for the recognition (by a Ub ligase) of specific degradation signals (degrons) within a target protein, resulting in the conjugation of Ub (usually in the form of polyUb) to a Lys residue within the target. Ultimately, the tagged protein is processed into short peptides by a single ATP-dependent protease (the 26S proteasome). The broad specificity of this system is achieved through the large number of Ub ligases (∼1,000 in mammals), which mediate recognition of different substrates. In contrast to the mammalian cytosol, protein degradation in eukaryotic organelles (similar to that in the bacterial cytosol) is mediated by several different ATP-dependent proteases, which function together with a handful of specialised adaptor proteins to enhance their substrate specificity ^16-18^. Collectively these machines, regardless of their subcellular localisation, are referred to as AAA+ (ATPases associated with a variety of cellular activities) proteases ^19-22^ as they are generally composed of two components, an unfoldase component belonging to the AAA+ superfamily and a peptidase component. In human mitochondria, five different AAA+ proteases have been discovered, two soluble matrix proteases; CLPXP (composed of two separate components; an unfoldase component CLPX and the peptidase component CLPP) and LONM (also known as LONP1) and three membrane bound proteases; an *intermembrane space-*AAA (*i*-AAA) protease and two forms of *matrix-*AAA (*m*-AAA) protease ^23^. Although LONM is considered to be the principal matrix quality control protease ^24-27^ there is growing evidence that CLPXP also plays a crucial role in mitochondrial proteostasis, contributing to heme regulation ^28-30^, mitoribosomal assembly ^31^ and the selective turnover of ROS-damaged subunits of Complex I ^32^. Consistently, mutations in CLPP that cause Perrault syndrome 3 are linked to mitochondrial dysfunction ^33-35^. Moreover, targeted dysregulation of mitochondrial CLPP was also recently demonstrated to be lethal to specific cancer cells ^36,37^. Despite the emerging importance of this protease complex in human mitochondria, our understanding of this machine and its mechanism of substrate recognition is largely based on homologous prokaryotic systems.

Here we show, that human PDIP38 is imported into isolated mitochondria, where it co-localizes with its partner proteins CLPX (and CLPP) in the matrix compartment. Importantly, PDIP38 does not trigger dissociation of the CLPXP complex, but rather we propose that PDIP38 is a mitochondrial adaptor protein for the AAA+ protease, CLPXP. Consistent with its role as a CLPX-adaptor protein, the structure of PDIP38 is composed of two domains, an N-terminal “YccV-like” domain which docks to an adaptor binding loop within the ZBD of CLPX, and a C-terminal DUF525-domain which forms an immunoglobulin-like fold bearing a putative substrate-binding groove. Significantly, the putative substrate-binding groove is lined with conserved hydrophobic residues and capped (at both ends) with conserved charged residues (basic at one end and acidic at the other). We speculate that, this groove is responsible for the specific recognition of short hydrophobic degrons, potentially located at the termini of a protein. Intriguingly, both domains have been found to recur throughout evolution (in bacteria, plants and humans), in select components of diverse protein degradation pathways. The YccV domain has been identified in bacterial (HspQ) and plant (ClpF) proteins, which appears to act as an anti-adaptor of the N-recognin, ClpS and hence regulates the recognition and turnover of N-degron bearing substrates ^38,39^. In contrast, the ApaG domain has been identified in Fbox-only proteins such as Fbx3, where it is proposed to act as a substrate recognition component. Consistent with this premise, we show that PDIP38 inhibits the LONM-mediated turnover of CLPX *in vitro* and stabilizes the steady state levels of CLPX *in vivo*. As such, PDIP38 regulates CLPXP activity, both directly (through the potential delivery of specific substrates to CLPX for degradation by CLPP) and indirectly (through inhibition of the LONM-mediated turnover of CLPX *in vivo*).

## RESULTS

### Human PDIP38 is a matrix localised mitochondrial protein

To date, mammalian PDIP38 has been identified in several subcellular compartments, from the plasma membrane to the nucleus and the mitochondrion ^1,6,12^, and as such its subcellular localisation is currently controversial. Therefore, to validate the sub-cellular localisation of human PDIP38, we performed *in vitro* import assays into isolated mitochondria and mitochondrial fractionation experiments (**Fig. 1**). Consistent with our identification of murine PDIP38 as a mitochondrial protein ^12^, radiolabelled human preprotein (pPDIP38) was imported into isolated mitochondria in a membrane potential-dependent manner (**Fig. 1a**, compare lanes 5 and 6). Importantly the processed, mature form of the protein (mPDIP38) was protected from cleavage by Proteinase K (Prot. K), demonstrating that mPDIP38 was sequestered inside the mitochondrion. Next, we performed a sub-mitochondrial fractionation of mitochondria isolated from HeLa cells coupled with a protease protection assays using Prot. K. Consistent with the location of PDIP38 within the mitochondrial matrix, and similar to a known matrix protein – CLPP, PDIP38 was protected from digestion by Prot. K in both intact mitochondria (**Fig. 1b**, lanes 1 – 4) and mitoplast (**Fig. 1b**, lanes 5 – 8). As a control, the outer membrane protein (TOM20) was digested by Prot. K under all conditions, while the inner membrane protein (TIM23) was completely protected in intact mitochondria but sensitive in mitoplast (**Fig. 1b**, lanes 5 – 8). Next, having established that human PDIP38 was indeed located within the mitochondrial matrix we analysed the interaction between CLPX and PDIP38 in human mitochondria by co-immunoprecipitation (co-IP). Initially, we examined the interaction of endogenous CLPX (with endogenous PDIP38), using a PDIP38 specific antisera immobilised to Protein A Sepharose (PAS). Consistent with a specific interaction between PDIP38 and CLPX, CLPX was only recovered in the presence of anti-PDIP38 (**Fig. 1c**, lane 3 lower panel) and not in the presence of the pre-immune sera (**Fig. 1c**, lane 2 lower panel). Next, to confirm this interaction we performed the reverse co-IP, in which the anti-CLPX antisera was immobilised to PAS. Consistent with the specific IP of CLPX with anti-PDIP38, the IP of CLPX using anti-CLPX antisera also resulted in the specific co-IP of PDIP38 (**Fig. 1d**, lane 3 lower panel).

**Figure 1.**
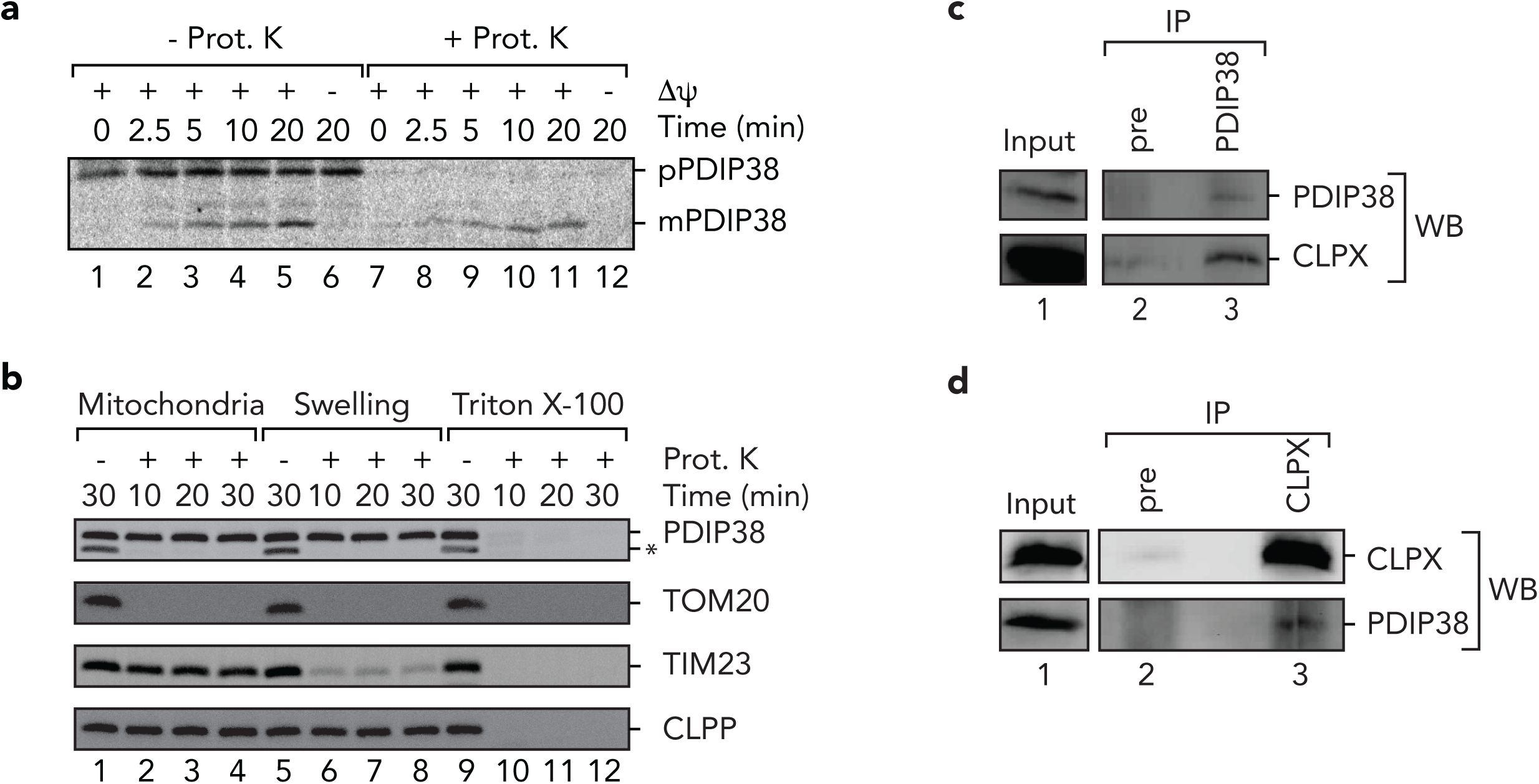
Human PDIP38 is imported into mitochondrial where it interacts with CLPX. **a.** Import of [^35^S]-labelled PDIP38 preprotein into mitochondria isolated from HeLa cells, in the presence or absence of a membrane potential (ΔΨ) as indicated, treated with (lanes 7 – 12) or without (lanes 1 – 6) proteinase K (Prot. K). Samples were separated by 15% Tris-glycine SDS-PAGE and analysed by digital autoradiography. **b.** Mitochondria were incubated, either in an osmotic buffer (lanes 1 – 4), isotonic buffer (Swelling) to rupture the outer membrane (lanes 5 – 8) or buffer containing Triton X-100 (lanes 9 – 12), in the absence (lanes 1, 5 and 9) or presence (lanes 2 – 4, 6 – 8 and 10 – 12) of Prot. K for the indicated time. Samples were separated by 15% Tris-glycine SDS-PAGE and subjected to immunoblotting with the appropriate antisera to visualize endogenous proteins. *, non-specific cross-reactive protein in PDIP38 antisera **c**. Specific immunoprecipitation of endogenous PDIP38 from detergent solubilized mitochondria using anti-PDIP38 polyclonal antibodies showing co-immunoprecipitation of endogenous CLPX. The input represents 50% of total mitochondrial lysate subjected to IP. **d**. Specific immunoprecipitation of endogenous CLPX from detergent solubilized mitochondria using anti-CLPX polyclonal antibodies showing co-immunoprecipitation of endogenous PDIP38. The input represents 50% of total mitochondrial lysate subjected to IP.

### The ZBD of CLPX is sufficient for interaction with PDIP38

Next, to ensure that the interaction observed *in vivo* (between PDIP38 and CLPX) was direct, we purified both components and examined the interaction *in vitro*. To do so, we generated a GST-PDIP38 fusion protein in which PDIP38 was fused to the C-terminus of GST. Following expression of GST-PDIP38, a soluble lysate (bearing overexpressed GST-PDIP38) was applied to Ni-NTA agarose beads (either lacking or containing immobilised His_10_-tagged CLPX (H_10_CLPX, see **Fig. 2a**). Following incubation of GST-PDIP38 with the beads, the specifically bound proteins were eluted from the column with imidazole (**Fig. 2b**, lanes 4 and 7). Consistent with our identification of PDIP38 as a novel CLPX interacting protein (**Fig. 1**), GST-PDIP38 was only recovered in the presence of immobilised H_10_CLPX (**Fig. 2b**, lanes 7) and not in the absence of an immobilised protein (**Fig. 2b**, lanes 4).

**Figure 2.**
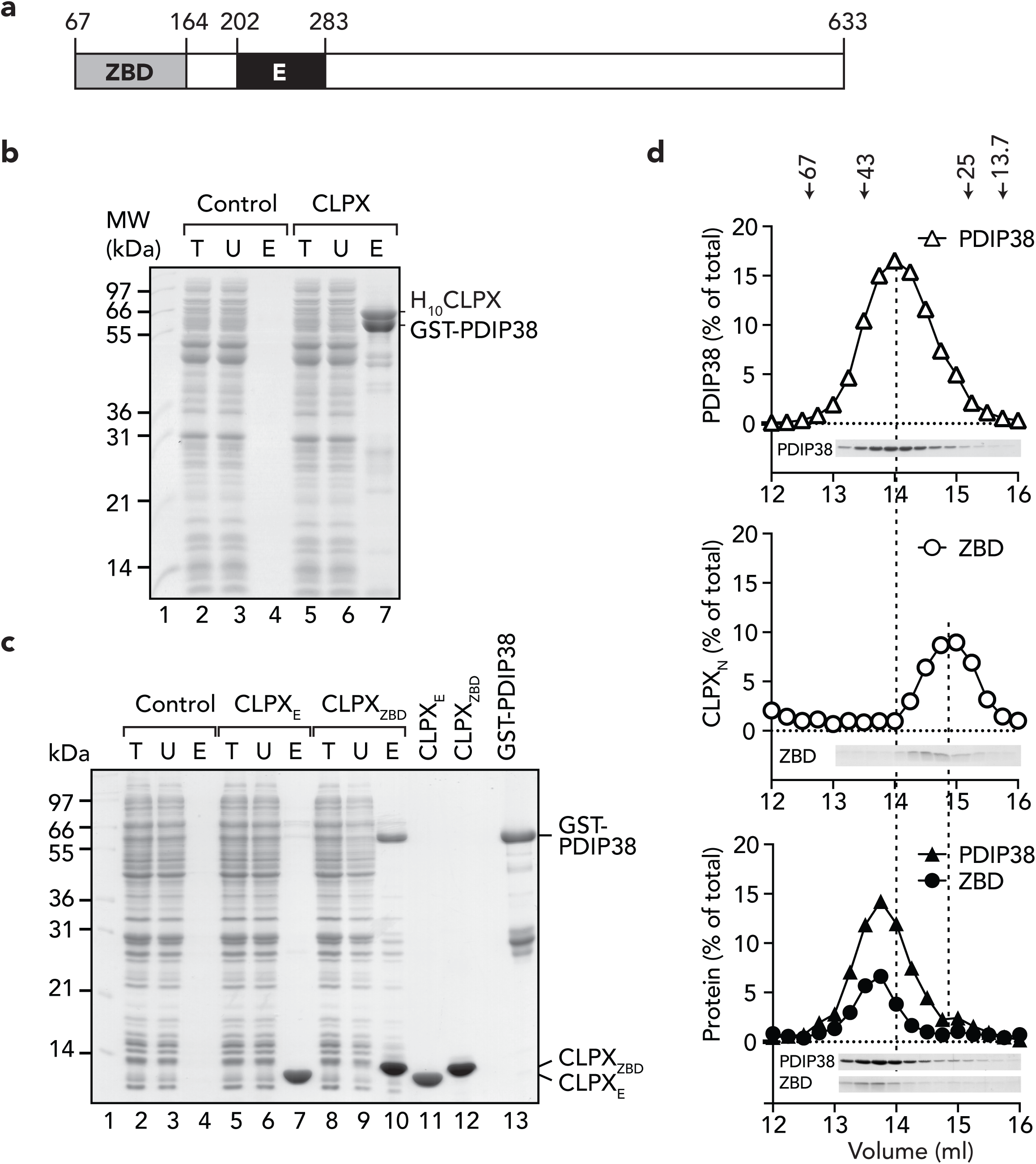
The interaction between PDIP38 and CLPX is mediated by the ZBD of CLPX. **a.** Cartoon representation of human CLPX domain structure. **b.** Coomassie stained 16.5% Tricine-buffered SDS-PAGE of PDIP38 pull-down from *E. coli* lysate expressing recombinant GST-PDIP38 using bead immobilised H_10_CLPX (CLPX) relative to beads only (control). T, total *E. coli* lysate with expressed GST-PDIP38; U, unbound fraction; E, eluted fraction. **c.** Coomassie-stained 16.5% Tricine-buffered SDS-PAGE of PDIP38 pull-down from *E. coli* lysate expressing recombinant GST-PDIP38 using bead immobilised H_10_CLPX E-domain (CLPX_E_) and H_10_CLPX Zinc binding domain (CLPX_ZBD_) relative to beads only (control). T, total lysate expressing GST-PDIP38; U, unbound fraction; E, eluted fraction. For comparison purified recombinant CLPX_E_ (lane 11), CLPX_ZBD_ (lane 12) and GST-PDIP38 (lane 13) are shown. **d.** Complex formation of PDIP38 and CLPX_ZBD_ was monitored by size exclusion chromatography (SEC) using a Superose 12 column. Elution profiles of PDIP38 (top panel), CLPX_ZBD_ (middle panel) or PDIP38 in the presence of CLPX_ZBD_ (bottom panel) were measured at 280 nm (A_280_). Arrows indicate the peak elution volume of Albumin (67 kDa), Ovalbumin (43 kDa), chymotrypsin A (25 kDa) and Ribonuclease (13.7 kDa).

Next, we asked the question how does the CLPX-PDIP38 complex form? Initially, we speculated that PDIP38 might be an adaptor protein of human CLPX, and as such would likely bind to an accessory domain of CLPX, as is the case for several bacterial AAA+ adaptor proteins ^16,40-42^. Interestingly in contrast to bacterial ClpX homologs, human CLPX contains two accessory domains, an N-terminal C4 type Zinc finger domain (often referred to as a Zinc binding domain (ZBD)) and an additional domain that is inserted into the AAA module of CLPX, which is unique to eukaryotic CLPX homologs (termed the E-domain ^43^). Therefore, to identify which CLPX domain (or domains) might be responsible for the interaction with PDIP38, we generated and purified H_10_-tagged versions of both the ZBD (**Fig. 2a**, ZBD) and the E-domain (**Fig. 2a**, E) of CLPX. These proteins were then immobilised to Ni-NTA agarose (as described above) and a soluble lysate bearing GST-PDIP38 was applied to the appropriate columns (**Fig. 2c**, lane 2, 5 and 8). Consistent with the data above, in which PDIP38 was recovered in the presence of full-length CLPX, PDIP38 was specifically co-eluted from the column containing immobilised ZBD (**Fig. 2c**, lane 10) and not from the column containing immobilised E-domain (**Fig. 2c**, lane 7). Collectively these data demonstrate that PDIP38 interacts specifically with the ZBD of CLPX. Next, we examined the stoichiometry of the interaction between PDIP38 and the N-domain. To address this question, we purified full-length PDIP38 (containing an N-terminal H_10_-tag) and monitored complex formation by size exclusion chromatography, using Superose 12. Both proteins alone formed a monodispersed peak on gel filtration, PDIP38 (alone) eluted in a single peak (**Fig. 2d**, upper panel), with an apparent molecular weight of ∼ 39 kDa (consistent with a monomeric protein in solution), while in contrast the ZBD of CLPX (**Fig. 2d**, middle panel) eluted at ∼15 ml, (which is consistent with an apparent molecular weight of ∼ 24 kDa and hence likely a homodimer of CLPX_ZBD_). Consistent with the pull-down (**Fig. 2d**), PDIP38 formed a stable complex with CLPX_ZBD_, which based on the elution volume of the complex (∼13.6 ml, equivalent to ∼ 49 kDa) likely forms a heterodimeric complex composed of one subunit of each protein (**Fig. 2d**, lower panel).

### PDIP38 is composed of two domains – only the NTD is required for interaction with CLPX

Having determined which domain of CLPX is required for interaction with PDIP38, we next examined the domain structure of PDIP38 with the aim of defining which region or regions in PDIP38 are required for interaction with the ZBD of CLPX. Initially we used a bioinformatic approach to determine the domain structure of PDIP38 ^44^. This analysis revealed that PDIP38 is composed of two domains, a large YccV-like N-terminal domain (residues 52 – 234) and a smaller C-terminal domain (residues 235 – 368) of unknown function (DUF525). Intriguingly, both domains have been identified in proteins that have been implicated in a variety of protein degradation pathways from bacteria to plants and humans. The YccV-like domain is present in the F-box protein Fbx21/FbxO21, which was recently demonstrated to form an integral component of the SCF (Skip-Cullin-Fbox) Ubiquitin (Ub) ligase complex required for the turnover of EID1 ^45,46^. This domain was also recently identified in a unique component (termed ClpF), which is proposed to form part of the Clp protease machinery in the chloroplast of *Arabidopsis thaliana* ^38^. In this case, ClpF was shown to form a binary complex with ClpS1, a putative adaptor protein (or N-recognin) that is required for the recognition of substrate proteins bearing specific N-degrons ^47,48^. Remarkably, bacterial YccV (recently renamed HspQ) was also shown to regulate protein turnover via two proteolytic system, activating Lon-mediated turnover and inhibiting ClpS-mediated degradation of N-degron substrates ^39,49^. Based on the bioinformatic analysis of PDIP38, we initially generated GST-fusion proteins of each domain, however unexpectedly neither fusion protein was soluble. Therefore, in order to identify a functional boundary of the proposed domains we performed limited proteolysis of mature PDIP38, using thermolysin (**Fig. 3a**). This approach revealed that PDIP38 was indeed composed of two stable domains (**Fig. 3a**, f1 and f2). However, based on the transient appearance of two intermediate fragments (f1’ and f2’), these domains are likely separated by an exposed, flexible linker. To identify the boundary of these two domains we performed six rounds of Edman degradation on fragment f1 (FLANHD). This defined f1 as the C-terminal DUF525 domain and identified the boundary of this domain as F157. Armed with this information we generated two additional GST-fusion proteins, GST-PDIP38_N_ (**Fig. 3b**) in which the N-terminal domain of PDIP38 (residues 52 to 153) was fused to the C-terminus of GST and GST-PDIP38_C_ (see **Fig. 3b**) in which the C-terminal domain of PDIP38 (residues 157 to 368) was fused to C-terminus of GST (GST-PDIP38_C_). To determine which domain was required for docking to CLPX we performed a series of pull-down assays, in which H_10_CLPX was immobilised to NiNTA-agarose beads and then incubated with a bacterial cell lysate containing either overexpressed GST-PDIP38, GST-PDIP38_N_ or GST-PDIP38_C_ (**Fig. 3c**, lanes 2, 4 and 6 respectively). As a control, the different GST-PDIP38 fusion proteins were also incubated with Ni-NTA-agarose beads lacking immobilised protein (**Fig. 3c**, lanes 3, 5 and 7, respectively). As expected, and consistent with Figure 2b, full length GST-PDIP38 was specifically eluted from the column containing immobilised H_10_CLPX (**Fig. 3c**, lanes 2). Significantly, deletion of the N-domain of PDIP38 (GST-PDIP38_C_) was sufficient to prevent any specific interaction between the two proteins (**Fig. 3c**, compare lanes 6 and 7). Consistent with these results, the N-domain of PDIP38 alone was sufficient for the interaction with CLPX as the GST-PDIP38_N_ fusion was specifically recovered from NiNTA-agarose beads containing immobilised CLPX (**Fig. 3c**, lanes 4) and not from beads lacking immobilised protein (**Fig. 3c**, lanes 5). Taken together these results demonstrate that the N-terminal YccV-like domain of PDIP38 specifically docks to the ZBD of CLPX.

**Figure 3.**
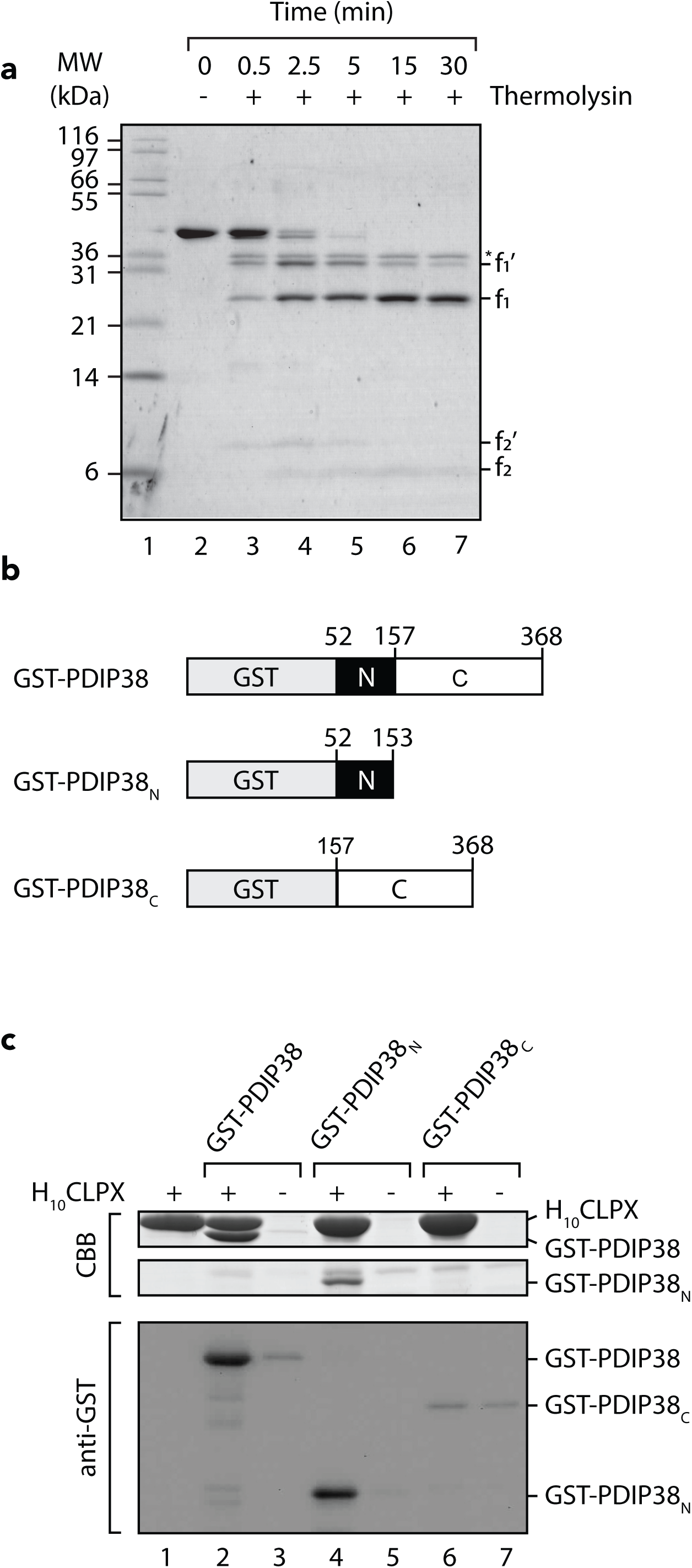
The N-terminal domain of PDIP38 interacts with CLPX. **a.** Limited proteolysis of native His_10_-tagged PDIP38 using thermolysin. Samples were analyzed by Coomassie stained 16.5% Tricine-buffered SDS-PAGE. *, thermolysin. f1, f1’, f2, f2’, fragments of PDIP38. **b.** Schematic representation of GST-PDIP38 fusion constructs. Preprotein numbering is used. **c.** *In vitro* pull-down using Ni-NTA agarose with (lanes 2, 4 and 6) or without (lanes 3, 5 and 7) purified immobilised H_10_CLPX, incubated with *E. coli* lysate expressing GST-PDIP38 (lanes 2 – 3), GST-PDIP38_N_ (lanes 4 – 5) or GST-PDIP38_C_ (lanes 6 – 7). Eluted fractions are shown with samples analyzed by Coomassie Brilliant Blue (CBB) staining or immunoblotting (with anti-GST) following separation by SDS-PAGE. As a control, purified H_10_CLPX (lane1) is shown.

### PDIP38 docks to the “adaptor binding” loop within the NTD of CLPX to regulate turnover of a model substrate

Next, we examined the consequence of PDIP38 docking to CLPX, i.e. is PDIP38 a substrate or an adaptor protein of CLPX, or does PDIP38 trigger dissociation of the CLPXP complex? To determine if PDIP38 is a substrate of CLPXP, we monitored the stability of PDIP38 *in vitro*, in the presence of active CLPXP (**Fig. 4a**). Given that substrate recognition by many Clp-proteases is generally mediated by degrons located at either the N- or C-termini ^50^, we generated an untagged version of PDIP38, using the Ub-fusion system ^51^. As a control, to ensure that human CLPXP was active, we monitored the turnover of the model unfolded protein (casein), a well-characterized CLPX substrate ^12,52^. Significantly, in contrast to the rapid CLPXP-mediated turnover of FITC-casein (**Fig. 4a**, middle panel), the levels of untagged PDIP38 remained unchanged throughout the time course of the experiment (**Fig. 4a**, upper panel). These data clearly demonstrate that mature untagged PDIP38 is not a substrate of the CLPXP protease, but rather is either an adaptor protein for human CLPX(P) or alternatively a protein “*switch*” that triggers dissociation of CLPP from the CLPXP complex. In order to address the second possibility and determine if PDIP38 is able to modulate the specificity of CLPXP, we next monitored the CLPXP-mediated turnover of FITC-labelled casein in the absence and presence of PDIP38 (**Fig. 4a**). Significantly, the addition of PDIP38 exhibited contrasting effects on the turnover of different forms of FITC-casein, specifically the CLPXP-mediated turnover of α_S2_casein was inhibited by PDIP38 (**Fig. 4a**, lower panel), in a concentration-dependent manner (**Supplementary Fig. 1)**, while the turnover of κ-casein was unaffected by the presence of PDIP38 (**Fig. 4a**, lower panel, **Supplementary Fig. 1**, black bars). Collectively these data suggest that PDIP38 was able to specifically inhibit the turnover of one substrate without affecting the turnover of another, demonstrating that PDIP38 does not trigger dissociation of CLPX from CLPP. In addition, PDIP38 itself was not a substrate of the CLPXP machine, suggesting that PDIP38 exhibits *adaptor-like* activity. To further investigate the possibility that PDIP38 is a CLPX adaptor protein we compared the ZBD of human CLPX with several other ClpX homologs, from both bacterial and eukaryotic species. In particular we focused on the known adaptor-docking region (**Fig. 4c**, adaptor binding loop). Despite considerable sequence conservation across the ZBD of bacterial and eukaryotic homologs, one region of the ZBD – the “*adaptor binding loop*” – diverged. This part of the domain was highly conserved amongst either eukaryotic (or bacterial) species but poorly conserved across the two kingdoms (**Fig. 4c and 4d**). Therefore, we hypothesized that this region may have coevolved with a new adaptor protein (i.e. PDIP38). To test the idea that the “*adaptor binding loop*” within the ZBD of human CLPX is required for docking to the putative adaptor protein PDIP38, we examined the ability of *Escherichia coli* ClpX (*ec*ClpX) ZBD (*ec*ZBD) to interact with human PDIP38 (**Fig. 4b**). Remarkably, in contrast to human ZBD (**Fig. 4b**, lane 1) which bound to PDIP38, *ec*ZBD failed to interact with PDIP38 at all (**Fig. 4b**, lane 3). Next to confirm that the proposed *adaptor binding loop* was indeed the site of PDIP38 interaction in human CLPX we replaced the putative adaptor-docking region (residues 120 to 123; SSTR) with AAAA in both full-length CLPX (here referred to as CLPX_4A_) and in the ZBD of CLPX (here referred to as ZBD_4A_). Consistent with the loss of binding of PDIP38 to ecZBD, the recovery of untagged PDIP38 to either immobilised ZBD_4A_ (**Fig. 4e**, lane 3) or CLPX_4A_ (**Fig. 4e**, lane 7) was completely abolished, when compared to wild type ZBD (**Fig. 4e**, lane 1) or CLPX (**Fig. 4e**, lane 5). Collectively these data suggest that the “*adaptor binding loop*” within the ZBD of CLPX performs a conserved function in both bacterial and eukaryotic homologs of CLPX. Specifically, the ZBD of human CLPX forms a crucial docking platform for interaction with the putative adaptor protein PDIP38. This interaction prevents the CLPXP-mediated turnover of the model substrate, α_S2_-casein, while permitting the turnover of κ-casein (**Fig. 4a**) suggesting that PDIP38 is a *bona fide* adaptor protein of mitochondrial CLPX.

**Figure 4.**
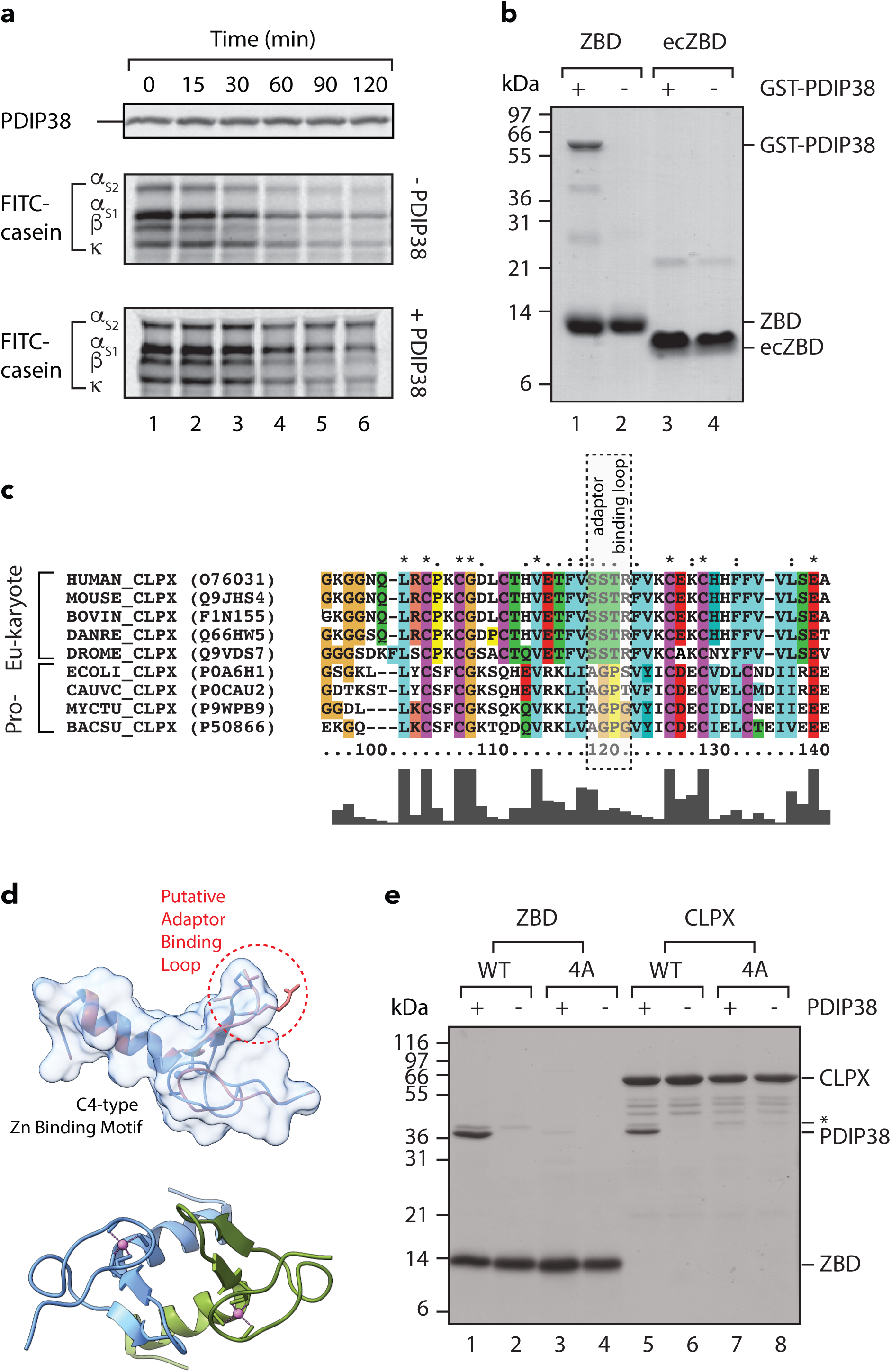
PDIP38 inhibits the CLPXP-mediated degradation of a model substrate via adaptor-like docking to CLPX_ZBD_. **a.** *In vitro* degradation of FITC-labelled casein by CLPXP protease in the absence and presence of 2.4 µM untagged PDIP38. Samples were separated by SDS-PAGE and analysed by fluorescence detection (FITC-casein) then CBB staining (PDIP38). **b.** *In vitro* pull-down, in which purified human (lanes 1 – 2) or *E. coli* (lanes 3 – 4) ZBD was immobilised to Ni-NTA agarose beads and incubated with (lanes 1 and 3) or without (lanes 2 and 4) an *E. coli* lysate expressing GST-PDIP38. Eluted fractions are shown with samples analysed by Coomassie Brilliant Blue (CBB) staining. **c.** Amino acid sequence alignment of eukaryotic and prokaryotic CLPX homologs showing the ZBD only. The *adaptor binding loop* in prokaryotic ClpX homologs is highlighted in the boxed section. ClpX sequences are *Homo sapiens* (O76031); *Mus musculus* (Q9JHS4); *Bovine* (F1N155); *Danio rerio* (Q66HW5), *Drosophila melanogaster* (Q9VDS7), *Escherichia coli* (P0A6H1), *Caulobacter crescentus* (P0CAU2), *Mycobacterium tuberculosis* (P9WPB9) and *Bacillus subtilus* (P50866). ***d.*** Ribbon diagram of *E. coli* CLPX_ZBD_ (PDB: 2DS6 ^58^, blue) overlaid with a model of the human CLPX_ZBD_ (red) highlighting the position of the putative adaptor-binding loop in human CLPX (circled). The space-filling model of *E. coli* ClpX N-domain is also shown in light blue. ***e.*** *In vitro* pull-down, in which purified wild type (lanes 1 – 2) or mutant (lanes 3 – 4) ZBD and wild type (lanes 5 – 6) or mutant (lanes 7 – 8) human H_10_CLPX was immobilised to Ni-NTA agarose beads and incubated with (lanes 1, 3, 5 and 7) or without (lanes 2, 4 and 6) an *E. coli* lysate expressing untagged PDIP38. Eluted proteins were separated by SDS-PAGE and visualized by Coomassie Brilliant Blue (CBB) staining.

Next, in order to determine the physiological function of PDIP38, we took two complementary approaches. In the first approach, we attempted to isolate PDIP38 interacting proteins. To do so, we knocked down PDIP38 expression in mammalian (HeLa) cells using siRNA with the aim of stabilising PDIP38-mediated substrates *in vivo*, before isolating the stabilised interacting proteins via pull-down. Although the knock down of PDIP38, using the PDIP38-specific siRNA (#22994, Thermo Fisher) was successful (**Fig. 5a**, middle panel compare lanes 1 – 3 with lanes 4 – 6), this approach was largely unproductive in identifying specific PDIP38 interacting proteins. Nevertheless, when analysing the steady state levels of selected mitochondrial proteins in the knock down cells, we noticed that the levels of CLPX were reduced in HeLa cells transfected with the PDIP38-specific siRNA (**Fig. 5a**, top panel, lanes 1 – 3) when compared to the levels of CLPX in HeLa cells transfected with a control siRNA (**Fig. 5a**, lanes 4 – 6). Importantly, this change was specific for CLPX as the levels of CLPP (**Fig. 5a**, lower panel) and the cross-reactive band (**Fig. 5a**, middle panel, *) were unchanged by knock down of PDIP38. To validate these data, we examined the steady state level of CLPX using two additional PDIP38-specific siRNA’s (s25055 and s25056) in comparison to an appropriate control siRNA (**Supplementary Fig. 2**). Significantly, the loss of CLPX (as a result of PDIP38 knock down) was specific, as the steady state levels of two unrelated proteins (i.e. mitochondrial SDHA or the cytosolic protein, GAPDH) were not affected (**Supplementary Fig. 2b**, lower panels).

**Figure 5.**
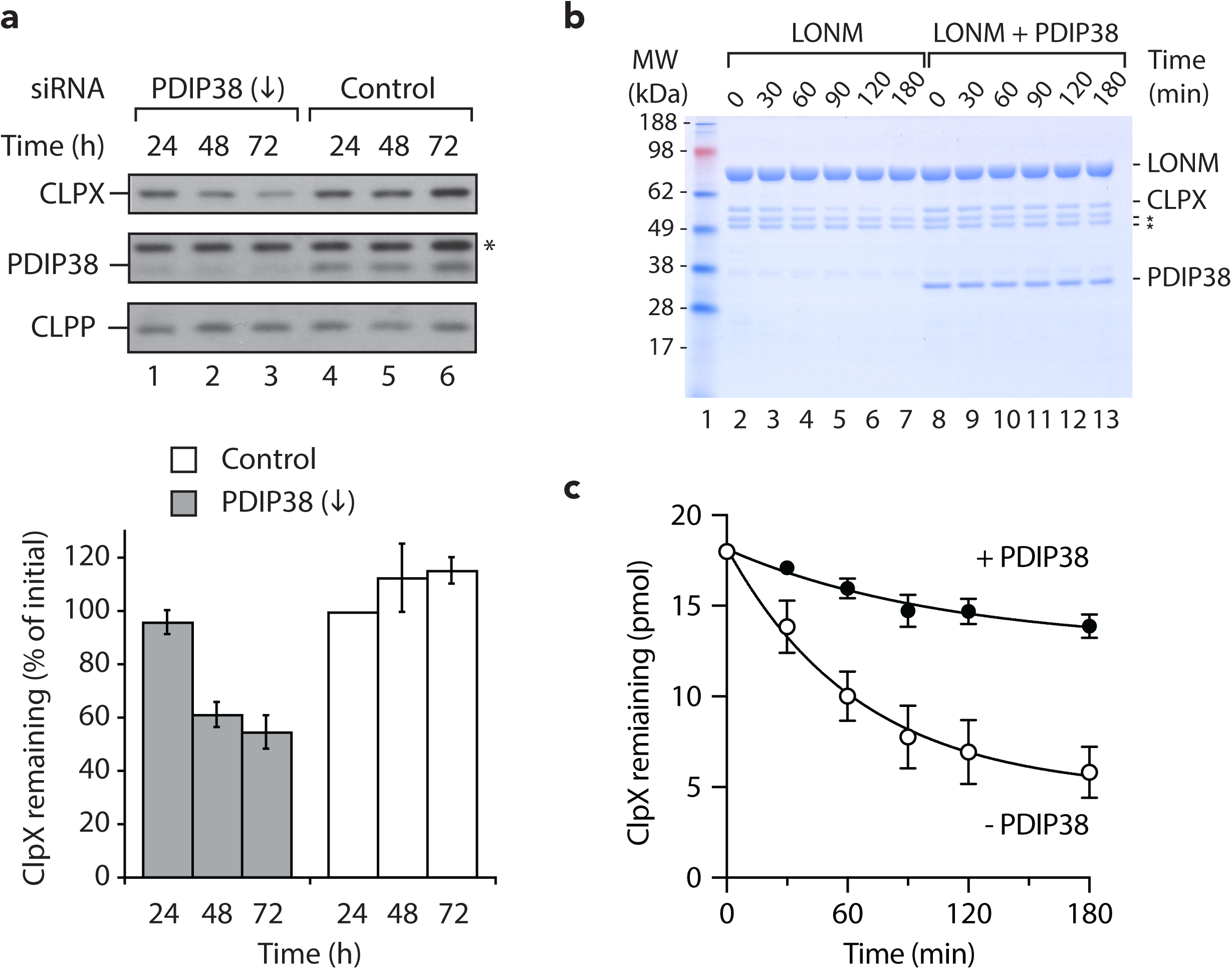
PDIP38 stabilises CLPX protecting it from LONM-mediated degradation. **a**. Representative immunoblots illustrating the steady state levels of CLPX (upper gel strip), PDIP38 (middle gel strip) and CLPP (lower gel strip) in PDIP38-depleted HeLa cells (lanes 1 – 3) relative to control HeLa cells (lanes 4 – 6). Samples were collected at the indicated times post-transfection of either Silencer Select siRNA (22994) targeting PDIP38 or a negative control siRNA (control). Proteins were separated by 16.5% Tris-Tricine SDS-PAGE. *, non-specific cross-reactive protein in PDIP38 antisera. The lower panel shows the quantitation of CLPX levels from three independent experiments, in PDIP38 depleted HeLa cells (grey bars) in comparison a negative control siRNA (white bars). Error bars represent the standard error of the mean (S.E.M.) of at least three independent experiments. **b.** *In vitro* degradation of CLPX by LONM_6_ protease (400 nM) in the absence or presence of 1 µM PDIP38. Samples were separated by 10% Tris-Tricine SDS-PAGE and analysed by CBB staining. * LONM impurity. **c.** Quantitation of *in vitro* degradation of CLPX by LONM_6_ (400 nM) in the absence (open symbols) or presence (closed symbols) of untagged PDIP38. Samples were separated by SDS-PAGE and analysed by CBB staining. Error bars represent the S.E.M. of three independent experiments.

From these data we speculated that PDIP38, similar to the *ec*ClpA adaptor protein *ec*ClpS (which protects its cognate unfoldase from auto-catalytic degradation *in vivo* ^40^), inhibits the autocatalytic turnover of CLPX. To test this, we monitored the stability of CLPX *in vitro*, in the presence of CLPP, with or without the addition of PDIP38. Contrary to the idea that PDIP38 inhibited auto-catalytic turnover of CLPX, the levels of CLPX (*in vitro*) remained unchanged in the presence of CLPP. Therefore, we hypothesized that the *in vivo* turnover of CLPX was mediated by an alternate mitochondrial matrix protease (i.e. LONM) and this turnover could be inhibited by PDIP38. To examine this possibility, we monitored the LONM-mediated degradation of CLPX *in vitro*, in the absence and presence of PDIP38 (**Fig. 5b**). Consistent with the idea that the levels of CLPX *in vivo* are controlled by the presence of PDIP38, CLPX was degraded by LONM *in vitro* (**Fig. 5b**, lanes 2 – 7) with a half-life of ∼ 60 min (**Fig. 5c**, open circles). Importantly, the LONM-mediated turnover of CLPX was inhibited by the addition of PDIP38 (**Fig. 5b**, lanes 8 – 13; **Fig. 5c**, filled circles). Significantly, the PDIP38-mediated inhibition of LONM was specific to the turnover of CLPX, as the LONM-mediated degradation of casein was unaffected by the addition of PDIP38 (**Supplementary Fig. 3**). Therefore, the inhibition of CLPX turnover is likely due to PDIP38 shielding the CLPX degron from interaction with LONM, which suggests that the CLPX degron is located within the ZBD of CLPX. Collectively, these data suggest that in the absence of PDIP38, the *in vivo* levels of CLPX may be regulated by LONM-mediated degradation.

Next, in order to better understand how substrate recognition by PDIP38 might occur, we crystalized mature PDIP38 (residues 52 to 368) and solved its structure by X-ray crystallography to 3.1 Å resolution (refer to Supplementary Table 1 for statistics). Consistent with our biochemical analysis (see **Fig. 3**), the structure of PDIP38 is composed of two domains, an N-terminal YccV-like domain (residues 64 to 186) and a C-terminal DUF525 domain (residues 231 to 368), separated by a short middle domain or linker region (**Fig. 6a**). The N-terminal YccV-like domain forms an anti-parallel β-sheet structure composed of six β-strands (β0-β5-β1-β2-β3-β4), in which strands β0 to β4 are connected by loops and β4 and β5 is connected by a short 3^10^ helix (**Fig. 6b** and **Supplementary Fig. 4**). In contrast to bacterial YccV (HspQ) homologs, PDIP38 contains a large insertion between β2 and β3, which forms an extended β-sheet that interacts with the proximal sheet of the DUF525 domain. Interestingly, this insertion is also present in other YccV-like proteins (including Human Fbx21), which lack the DUF525 domain. However, similar to PDIP38, Fbx21 contains an additional domain that is proposed to be involved in substrate-binding. Hence, we propose that similar to PDIP38, the extended β2/β3 sheet in Fbx21 is likely involved in a stabilising interaction with an associated substrate binding domain. In addition to the extension of the β2/β3 strands, PDIP38 also contains a unique insertion located between β3 and β4 (residues 143-166). This insertion is not only exposed (as it was susceptible to partial proteolysis) but is also highly flexible as it was not visible in the structure, presumably due to disorder. Based on the expected location of this loop, suspended over the DUF525 domain, we speculate that the L4 loop regulates substrate binding to the C-terminal domain. The linker region (or middle domain), which connects the N- and C-terminal domains, is formed by a small N-terminal α-helix (α1), a two-stranded anti-parallel β-sheet (β6 and β7) and a C-terminal α-helix (α2). This domain makes extensive contact to the N-terminal domain, wrapping around the domain, and likely forms a flexible hinge point for movement of the C-terminal domain and hence delivery of bound cargo to the associated ATPase component, CLPX. The C-terminal DUF525 domain (residues 231 to 368) forms an Immunoglobin-like β-sandwich fold composed of two four-stranded antiparallel β-sheets. The proximal sheet is composed of β8-β9-β10-β13, while the distal sheet is composed of strands β12-β11-β14-β15. Interestingly, although this domain exhibits only limited amino acid identity (∼30%) with bacterial ApaG proteins and eukaryotic F-box only proteins such as Fbx3, this group of proteins share considerable structural homology (root-mean-squared deviation (RMSD) of ∼1.5 A. Consistent with our identification of PDIP38 as a putative substrate delivery factor for mitochondrial CLPX, Fbx3 forms part of a SCF Ubiquitin ligase complex in which the DUF525 domain is proposed to be involved in substrate recognition ^53,54^. Therefore, to gain further insight into the function PDIP38 we examined the molecular surface of our structure (**Fig. 6c**). From this analysis we identified a conserved hydrophobic groove, located between the two β-sheets of the C-terminal domain, which is flanked by conserved charged residues at opposite ends of the groove (**Fig. 6d**). To determine the significance of this groove we also examined the surface of *Xanthomonas axonopodis* ApaG (PDB: 2F1E) and human Fbx3 (PDB:5HDW). Significantly, despite the weak overall sequence similarity across this group of proteins, the physicochemical properties of this groove are remarkably conserved across all three proteins (**Fig. 6e and 6f, Supplementary Fig. 5**). Indeed, all but one of the 9 hydrophobic residues that line the hydrophobic pocket and both of the charged residues that flank the groove are absolutely conserved from bacteria to humans (see **Supplementary Table 2 and Supplementary Fig. 5b**). Furthermore, of the absolutely conserved residues that found within this domain, approximately half of them are clustered to the hydrophobic groove. Consistent with the notion that this conserved groove plays an important role in substrate recognition, Chen and colleagues discovered a small molecule inhibitor of Fbx3, that docks into the conserved hydrophobic pocket where it is proposed to make a crucial interaction with the conserved acid residue that caps the groove ^55^.

**Figure 6.**
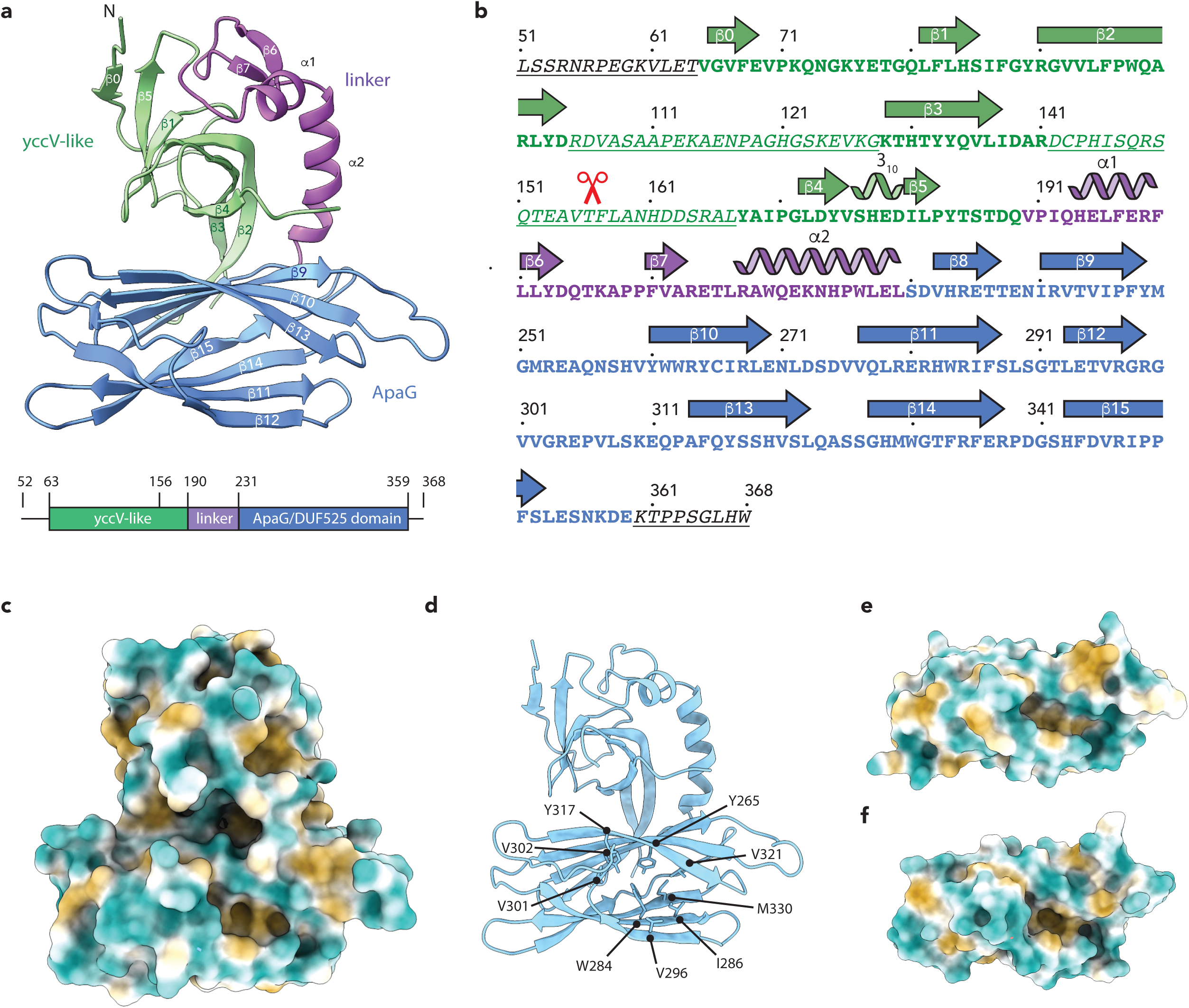
Structure of PDIP38 highlighting the conserved hydrophobic groove. **a**. Ribbon representation of human PDIP38 highlighting its three domains. The N-terminal yccV-like domain (green) and the C-terminal DUF525 domain (blue) are separated by a hinge or linker domain (magenta). **b.** Secondary elements are illustrated above the amino acid sequence. Red scissors indicate the site of cleavage by thermolysin. Underlined protein sequence represents disordered regions of the protein structure. **c.** Ribbon representation of human PDIP38, highlighting the conserved hydrophobic residues that line the binding groove. **d-f**. Hydrophobic surface representation of (d) PDIP38 illustrating the substrate binding groove compared to (e) Human Fbx3 DUF525 domain (PDB code 5HDW) and (f) *Xanthomonas axonopodis* ApaG (PDB code 2F1E). All figures were generated in ChimeraX_Daily.

What type of protein substrates might PDIP38 recognise? One possibility, that would be specific to the matrix compartment, is the recognition of incompletely or aberrantly processed matrix proteins, which retain their N-terminal presequence. These presequences are generally enriched in hydrophobic and basic residues, hence we speculate that PDIP38 might be responsible for the recognition these incorrectly processed or unprocessed matrix proteins, delivering them to CLPXP for removal. This role is somewhat similar to that of the *E. coli* ClpX adaptor protein – SspB (which is responsible for the recognition of incomplete translation products that bear a largely hydrophobic C-terminal recognition motif known as the SsrA-tag ^41,56,57^. An alternate possibility involves the more general recognition of exposed hydrophobic patches that are found in misfolded proteins that accumulate in response proteostatic stress. Conversely, PDIP38 may be required for the conditional recognition of proteins that expose hydrophobic motifs at either their N- or C-termini.

In summary, we show that PDIP38 is a novel component of the proteostasis network in mammalian mitochondria. Not only does PDIP38 modulate CLPXP substrate specificity and inhibit the LONM-mediated turnover of CLPX *in vitro*, but it also stabilises the steady state levels of CLPX *in vivo*. As such we propose that PDIP38 represents a novel mitochondrial adaptor protein for the CLPXP protease. Consistently, the atomic structure of PDIP38 revealed that the protein is composed of two domains separated by an α-helical hinge. The N-terminal YccV-like domain is crucial for interaction with the adaptor binding loop within the ZBD of CLPX, while the C-terminal domain contains a conserved hydrophobic groove which is proposed to facilitate substrate binding and hence delivery to CLPX. Significantly, the residues that line this hydrophobic groove are highly conserved across DUF525 containing proteins, from bacteria to humans. Hence, we speculate that the bacterial PDIP38 homolog (i.e. ApaG) may also play a role in protein turnover. An important challenge for the future will be the *in vivo* dissection of this system to identify the physiological substrates of the CLPXP protease that are delivered by PDIP38.

## ACKNOWLEDGEMENTS

This work was supported by an Australian Research Council (ARC) Discovery Project (DP0770013) to D.A.D and K.N.T., and ARC Future Fellowship to K.N.T. (FT0992033) and an ARC Australian Research Fellowship to D.A.D (DP110103936). P.R.S. and H.Z were supported by a La Trobe University Postgraduate Award, E.J.B. was supported by Australian Postgraduate Awards and T.S. was supported Cooperative Research Centre postgraduate award. We thank Mia Miasari for cloning of *PDIP38*_*N*_ and *PDIP38*_*C*_ into pGEX-4T.

## Author contribution

Conceptualization, D.A.D. and K.N.T.; Methodology, D.A.D., K.N.T. and K.Z.; Investigation, P.R.S., E.J.B., H.Z., V.J.S., L.J.V., T.S. and K.Z.; Writing – Original Draft, D.A.D., K.N.T. and K.Z.; Supervision, Project Administration and Funding Acquisition, D.A.D. and K.N.T.

## Methods

### Plasmids

For *in vitro* transcription and translation of human PDIP38, pOTB7/*PDIP38* was obtained from the I.M.A.G.E. Consortium (ID 3349399). For the heterologous expression of PDIP38 in *E. coli*, the cDNA coding for mature PDIP38 (residues 52-368) was amplified by PCR from pOTB7/*PDIP38* using the appropriate primers (Supplementary Table 3) and cloned into either pHUE ^51^ between *Sac* II and *Hind* III (to express untagged PDIP38), pET10N ^59^ between *Not* I and *Xho* I (to express PDIP38 with an N-terminal H_10_ tag), pET10C ^59^ between *Nde* I and *Not* I (to express PDIP38 as a C-terminal H_10_ fusion protein), or pGEX-4T-1 between *Bam* HI and *Xho* I (to express PDIP38 as an N-terminal Glutathione S-transferase (GST) fusion protein). To generate PDIP38_N_ (residues 52-153) and PDIP38_C_ (residues 157-368) fused to GST, pGEX-4T/*PDIP38* was subjected to site-directed mutagenesis ^60^ using primers PDIP_bam1 and PDIP_bam2 (see Supplementary Table 3). The resulting plasmid (pDT1367, see Supplementary Table 4) contained a stop codon and an additional *Bam* HI site (and was used directly for expression of GST-PDIP38_N_). To generate GST-PDIP38_C_, pDT1367 was digested with *Bam* HI, the cut vector ligated lacking the *PDIP38*_*N*_ fragment to generate pDT1362. Plasmids for bacterial expression of human CLPX (full-length and domain mutants) and human CLPP (either His-tagged and untagged) were described previously ^33^. For expression of CLPX_4A_ and ZBD_4A_, pET10C/*hCLPX*_*4A*_ and pET10C/*hZBD*_*4A*_ were generated by site directed mutagenesis using appropriate primers (see Supplementary Table 3). For details of primer sequences and plasmid constructs, refer to Supplementary information. All clones were confirmed by Sanger sequencing.

### Proteins

Recombinant proteins were expressed, either in BL21-CodonPlus®(DE3)-RIL or XL1-Blue (Agilent) *Escherichia coli* cells, as appropriate. His-tagged (H_6_- or H_10_-) recombinant proteins were purified from clarified *E. coli* lysates under native conditions by immobilised metal affinity chromatography using Ni-NTA agarose (Qiagen) essentially as described ^12^ using 50 mM Tris-HCl, [pH 8.0], 300 mM NaCl supplemented with an appropriate concentration of imidazole for binding (10 or 20 mM), washing (20 mM or 65 mM) and elution (250 mM or 500 mM). Purified His_6_-Ub-PDIP38 and His_6_-Ub-CLPP were cleaved using His_6_-Usp2cc ^51^ and the untagged mature proteins recovered via a method outlined previously ^51,61^. GST-PDIP38 was purified by affinity chromatography using GSH agarose (Bioserve) as outlined by the manufacturer. Radiolabelled PDIP38 preprotein was synthesized using TNT® SP6 Quick Coupled Transcription-Translation System (Promega) with undigested pOTB7/*PDIP38* as template and 11 μCi of [^35^S]Met/CysEXPRE^35^S^35^S protein labelling mix (specific activity of >1000 Ci/mmol) from Perkin Elmer. Protein Assay (Bio-Rad) was used to determine protein concentrations using bovine serum albumin (Thermo Scientific) as a standard. Protein concentrations refer to the protomer unless otherwise stated. FITC-casein, thermolysin, proteinase K and lysozyme were purchased from Sigma-Aldrich, DNase I was purchased from Gold Biotechnology. SeeBlue® Plus2 pre-stained and Mark12™ unstained protein standards were from Life Technologies.

### Electrophoresis and protein detection

Proteins were separated using either glycine- or Tricine-buffered ^62^ SDS-PAGE. Protein samples in 1 x SDS-PAGE sample buffer (80 mM Tris-HCl [pH 6.8], 2% (w/v) SDS, 5% (v/v) glycerol, 100 mM DTT and 0.02% (w/v) bromophenol blue) were heat treated at 95 °C for 5 min before separation. For visualization of proteins, gels were stained with Coomassie Brilliant Blue R250 solution (CBB) or transferred to polyvinyldiflouride (PVDF) membrane using semi-dry method for immunoblotting. Primary antibodies used were anti-PDIP38 (POLDIP2; Abcam), anti-PDIP38 (125/88; generated in rabbit using purified recombinant PDIP38-H_10_ as antigen), affinity purified anti-CLPX ^12^, anti-LONM ^12^, anti-TIM23 (BD Biosciences), anti-SDHA (Invitrogen), anti-GST (GE Healthcare), anti-GAPDH (Life Technologies) and anti-mtHSP60 (N. Hoogenraad, La Trobe University). Peroxidase coupled secondary antibodies were anti-rabbit, anti-mouse and anti-goat IgG (Sigma-Aldrich). Antibody complexes were detected using enhanced chemiluminescence detection reagents (GE Healthcare) and digital images captured using GeneSnap (SynGene) or Image Lab™ (Bio-Rad) Software. FITC-casein was detected by in gel fluorescence (excitation 488 nm and emission 526 nm) while radiolabelled proteins were detected by exposing dried gels to phosphor screens. Imaging was performed using a Typhoon™ Trio variable mode imager and analysed using ImageQuant software (GE Healthcare).

### Limited proteolysis

H_10_PDIP38 (0.1 mg/ml) was subjected to limited proteolysis using thermolysin (0.01 mg/ml) at 30°C in 50 mM Tris-HCl [pH 7.0], 150 mM NaCl and 5 mM CaCl_2_. To terminate the reaction, samples were treated with 2 mM PMSF and heated at 95 °C in 1 x SDS-PAGE sample buffer.

### Degradation assays

The CLPXP-mediated degradation of FITC-casein was performed essentially as described ^52^. Briefly, 0.4 µM CLPX_6_P_14_ was preincubated (at 30 °C for 5 min) in proteolysis buffer (50 mM Tris-HCl [pH 8.0], 100 mM KCl, 20 mM MgCl_2,_ 1 mM DTT, 0.02 % (v/v) Triton X-100, 10 % (v/v) glycerol) with FITC-casein (0.3 µM) in the absence or presence of 2.4 µM untagged PDIP38. To initiate degradation 5 mM ATP was added and samples were incubated at 30 °C for the times indicated. Reactions were terminated by the addition of 1 x sample buffer and the proteins denatured at 95 °C for 5 min.

### In vitro binding analysis

The *in vitro* binding analysis was adapted from the method outlined in ^63^. *E. coli* cells containing expressed GST-PDIP38, GST-PDIP38_N_, GST-PDIP38_C_ or untagged PDIP38, were resuspended (5 ml/g wet weight of cells) in Binding Buffer (20 mM HEPES-KOH [pH 7.5], 100 mM K(OAc), 10 mM Mg(OAc), 10 % (v/v) glycerol, 65 mM imidazole) supplemented with 0.5 % (v/v) Triton X-100, EDTA free protease inhibitor cocktail (Roche), 2 mM PMSF and DNase I (10 μg/ml) then subjected to chemical lysis with lysozyme (0.2 mg/ml). Cell free lysates or purified untagged PDIP38, as appropriate, were applied to Ni-NTA agarose beads either lacking or containing immobilised H_10_-tagged CLPX, CLPX_ZBD_, CLPX_E,_ *ec*ClpX_ZBD_, CLPX_4A_ or ZBD_4A_ and incubated with end-over-end mixing at 4 °C for 30 min. The beads were then washed with 5 bed volumes (BV) of Binding Buffer supplemented with 0.5 % (v/v) Triton X-100 followed by 10 BV of Wash Buffer (Binding buffer supplemented with 0.25 % (v/v) Triton X-100). Bound proteins were eluted with Elution Buffer (50 mM Tris-HCl [pH 8.0], 300 mM NaCl, 500 mM imidazole). For binding assays containing full length wild type or mutant CLPX, all buffers were supplemented with 2 mM ATP and 10 mM β-mercaptoethanol.

### Cell culturing and treatment

HeLa cells were cultured in Dulbecco’s Modified Eagle’s Medium (Life Technologies) supplemented with 10 % (v/v) fetal calf serum at 37 °C under an atmosphere of 5 % (v/v) CO_2_. Transfection of plasmid (10 µg) or 10-20 nM synthetic siRNA (Life Technologies) was performed using Lipofectamine® 2000 transfection reagent (Life Technologies) as per the manufacturer’s instructions and cells grown for a further 24-72 h, as indicated. For interference of *PDIP38* mRNA three independent synthetic siRNA (Life Technologies) were used; Silencer No. 22994 and Silencer Select No. s25055 (s55) and s25056 (s56). The corresponding Silencer Negative Control and Silencer Select Negative Controls No. 1 (nc1) and No. 2 (nc2) were used. For analysis, cells where detached by trypsin treatment (0.25% (w/v) trypsin, 1mM EDTA; Invitrogen) and washed cell pellets lysed using TC extraction buffer (50 mM Tris-HCl [pH 7.5], 375 mM NaCl, 1 mM EDTA, 1% (v/v) Triton X-100) freshly supplemented with 2 mM phenylmethanesulfonyl fluoride (PMSF). Soluble lysate was collected and used for analysis.

### Mitochondrial isolation and manipulation

Crude mitochondria were isolated from HeLa cells as described ^12,64^. *In vitro* import ^65^ was performed at 37 °C with [^35^S]Met/Cys-labelled preprotein and isolated mitochondria resuspended in Import Buffer (20 mM HEPES-KOH [pH 7.4], 250 mM sucrose, 5 mM Mg(OAc), 80 mM K(OAc), freshly supplemented with 10 mM Na succinate, 1 mM DTT, 2% (w/v) fatty acid free BSA, 5 mM ATP and 5 mM methionine. A mix of valinomycin (2 µM) and oligomycin (10 µM) was used to dissipate the membrane potential. Following import, mitochondria resuspended in SEM (250 mM sucrose, 1 mM EDTA, 10 mM MOPS-KOH [pH 7.2]) were treated with ∼40 µg/ml proteinase K (Prot. K) for 15 min at 4 °C. Mitoplasts were formed in 9 parts EM buffer (10 mM MOPS-KOH [pH 7.2], 1 mM EDTA) to 1 part SEM buffer at 4 °C for 20 min with gentle pipetting ^65^. For protease treatment, mitochondria in SEM buffer, mitoplasts in EM buffer and lysed mitochondria in SEM buffer with 0.5 % (v/v) Triton X100 were incubated on ice with 50 μg/ml proteinase K for the times indicated. Proteinase K was inhibited by the addition of 2 mM phenylmethylsulfonyl fluoride (PMSF) and proteins were immediately precipitated with TCA for analysis.

### Immunoprecipitation

Mitochondrial lysate in immunoprecipitation (IP) Buffer (50 mM Tris-HCl [pH 7.5], 100 mM KCl, 10 mM Mg(OAc), 5% v/v glycerol) containing 0.5 % (v/v) Triton X-100, 10 mM ATP and 2 mM phenylmethylsulfonyl fluoride (PMSF) was mixed with Protein A-Sepharose covalently attached to antibodies (anti-PDIP38 or anti-CLPX) by end-over-end rotation for 1 h, at 4°C. Beads were washed with 3 bed volume (BV) of IP buffer containing 0.25 % (v/v) Triton X-100, 10 mM ATP and 2 mM PMSF and antibody bound protein eluted using 1 BV of 50 mM glycine [pH 2.5].

### Crystallisation, X-ray diffraction and structure determination

To investigate the structure of PDIP38, crystal screening was performed and crystals were obtained using 20% (w/v) PEG8000, 100 mM Hepes, pH 7.5. Crystals of the free and derivatized protein were frozen in liquid nitrogen and data were collected at 100 K at the Swiss light source (SLS, Villigen, Switzerland; beamline PXII). Data were recorded on a PILATUS 6M detector (Dectris, Baden-Daettwil, Switzerland) and data reduction was performed using the program package XDS ^66,67^. The structure of PDIP38 was solved to 3.1 Å by single anomalous dispersion techniques using one Pt derivative for phasing. The model was refined using PHENIX ^68^. Most of the structure was unambiguously assigned in the electron density map except for residues 52–62 at the N-terminus and the loop regions (L3 between residues 108–126) and (L4 between residues 144–167), due to poor density. Supplementary Table 1 provides the statistics for the X-ray data collection and final refined model. All structural figures were generated using ChimeraX_Daily.

## References

1 Liu, L., Rodriguez-Belmonte, E. M., Mazloum, N., Xie, B. & Lee, M. Y. Identification of a novel protein, PDIP38, that interacts with the p50 subunit of DNA polymerase delta and proliferating cell nuclear antigen. J Biol Chem 278, 10041–10047, doi:10.1074/jbc.M208694200 (2003).

2 Klaile, E., Kukalev, A., Obrink, B. & Muller, M. M. PDIP38 is a novel mitotic spindle-associated protein that affects spindle organization and chromosome segregation. Cell Cycle 7, 3180–3186, doi:10.4161/cc.7.20.6813 (2008).

3 Tissier, A. et al. Crosstalk between replicative and translesional DNA polymerases: PDIP38 interacts directly with Poleta. DNA Repair (Amst) 9, 922–928, doi:10.1016/j.dnarep.2010.04.010 (2010).

4 Wong, A. et al. PDIP38 is translocated to the spliceosomes/nuclear speckles in response to UV-induced DNA damage and is required for UV-induced alternative splicing of MDM2. Cell Cycle 12, 3184–3193, doi:10.4161/cc.26221 (2013).

5 Xie, B. et al. Further characterization of human DNA polymerase delta interacting protein 38. J Biol Chem 280, 22375–22384, doi:10.1074/jbc.M414597200 (2005).

6 Klaile, E. et al. The cell adhesion receptor carcinoembryonic antigen-related cell adhesion molecule 1 regulates nucleocytoplasmic trafficking of DNA polymerase delta-interacting protein 38. J Biol Chem 282, 26629–26640, doi:10.1074/jbc.M701807200 (2007).

7 Arakaki, N. et al. Regulation of mitochondrial morphology and cell survival by Mitogenin I and mitochondrial single-stranded DNA binding protein. Biochim Biophys Acta 1760, 1364–1372, doi:10.1016/j.bbagen.2006.05.012 (2006).

8 Sutliff, R. L. et al. Polymerase delta interacting protein 2 sustains vascular structure and function. Arterioscler Thromb Vasc Biol 33, 2154–2161, doi:10.1161/ATVBAHA.113.301913 (2013).

9 Lyle, A. N. et al. Poldip2, a novel regulator of Nox4 and cytoskeletal integrity in vascular smooth muscle cells. Circ Res 105, 249–259, doi:10.1161/CIRCRESAHA.109.193722 (2009).

10 Kim, Y. et al. Essential role of POLDIP2 in Tau aggregation and neurotoxicity via autophagy/proteasome inhibition. Biochem Biophys Res Commun 462, 112–118, doi:10.1016/j.bbrc.2015.04.084 (2015).

11 Grinchuk, O. V., Motakis, E. & Kuznetsov, V. A. Complex sense-antisense architecture of TNFAIP1/POLDIP2 on 17q11.2 represents a novel transcriptional structural-functional gene module involved in breast cancer progression. BMC Genomics 11 Suppl 1, S9, doi:10.1186/1471-2164-11-S1-S9 (2010).

12 Lowth, B. R. et al. Substrate recognition and processing by a Walker B mutant of the human mitochondrial AAA+ protein CLPX. J Struct Biol 179, 193–201, doi:10.1016/j.jsb.2012.06.001 (2012).

13 Cheng, X. et al. PDIP38 associates with proteins constituting the mitochondrial DNA nucleoid. J Biochem 138, 673–678, doi:10.1093/jb/mvi169 (2005).

14 Paredes, F. et al. Poldip2 is an oxygen-sensitive protein that controls PDH and alphaKGDH lipoylation and activation to support metabolic adaptation in hypoxia and cancer. Proc Natl Acad Sci U S A 115, 1789–1794, doi:10.1073/pnas.1720693115 (2018).

15 Sala, A. J., Bott, L. C. & Morimoto, R. I. Shaping proteostasis at the cellular, tissue, and organismal level. J Cell Biol 216, 1231–1241, doi:10.1083/jcb.201612111 (2017).

16 Kirstein, J., Moliere, N., Dougan, D. A. & Turgay, K. Adapting the machine: adaptor proteins for Hsp100/Clp and AAA+ proteases. Nat Rev Microbiol 7, 589–599, doi:10.1038/nrmicro2185 (2009).

17 Nishimura, K. & van Wijk, K. J. Organization, function and substrates of the essential Clp protease system in plastids. Biochim Biophys Acta 1847, 915–930, doi:10.1016/j.bbabio.2014.11.012 (2015).

18 Varshavsky, A. The N-end rule pathway and regulation by proteolysis. Protein Sci 20, 1298–1345, doi:10.1002/pro.666 (2011).

19 Gur, E., Ottofueling, R. & Dougan, D. A. Machines of destruction - AAA+ proteases and the adaptors that control them. Subcell Biochem 66, 3–33, doi:10.1007/978-94-007-5940-4_1 (2013).

20 Olivares, A. O., Baker, T. A. & Sauer, R. T. Mechanical Protein Unfolding and Degradation. Annu Rev Physiol 80, 413–429, doi:10.1146/annurev-physiol-021317-121303 (2018).

21 Striebel, F., Kress, W. & Weber-Ban, E. Controlled destruction: AAA+ ATPases in protein degradation from bacteria to eukaryotes. Curr Opin Struct Biol 19, 209–217, doi:10.1016/j.sbi.2009.02.006 (2009).

22 Yu, H. & Matouschek, A. Recognition of Client Proteins by the Proteasome. Annu Rev Biophys 46, 149–173, doi:10.1146/annurev-biophys-070816-033719 (2017).

23 Glynn, S. E. Multifunctional Mitochondrial AAA Proteases. Front Mol Biosci 4, 34, doi:10.3389/fmolb.2017.00034 (2017).

24 Bezawork-Geleta, A., Brodie, E. J., Dougan, D. A. & Truscott, K. N. LON is the master protease that protects against protein aggregation in human mitochondria through direct degradation of misfolded proteins. Sci Rep 5, 17397, doi:10.1038/srep17397 (2015).

25 Bota, D. A. & Davies, K. J. Lon protease preferentially degrades oxidized mitochondrial aconitase by an ATP-stimulated mechanism. Nat Cell Biol 4, 674–680, doi:10.1038/ncb836 (2002).

26 Fukuda, R. et al. HIF-1 regulates cytochrome oxidase subunits to optimize efficiency of respiration in hypoxic cells. Cell 129, 111–122, doi:10.1016/j.cell.2007.01.047 (2007).

27 Tian, Q. et al. Lon peptidase 1 (LONP1)-dependent breakdown of mitochondrial 5-aminolevulinic acid synthase protein by heme in human liver cells. J Biol Chem 286, 26424–26430, doi:10.1074/jbc.M110.215772 (2011).

28 Kubota, Y. et al. Novel Mechanisms for Heme-dependent Degradation of ALAS1 Protein as a Component of Negative Feedback Regulation of Heme Biosynthesis. J Biol Chem 291, 20516–20529, doi:10.1074/jbc.M116.719161 (2016).

29 Yien, Y. Y. et al. Mutation in human CLPX elevates levels of delta-aminolevulinate synthase and protoporphyrin IX to promote erythropoietic protoporphyria. Proc Natl Acad Sci U S A 114, E8045–E8052, doi:10.1073/pnas.1700632114 (2017).

30 Kardon, J. R. et al. Mitochondrial ClpX Activates a Key Enzyme for Heme Biosynthesis and Erythropoiesis. Cell 161, 858–867, doi:10.1016/j.cell.2015.04.017 (2015).

31 Szczepanowska, K. et al. CLPP coordinates mitoribosomal assembly through the regulation of ERAL1 levels. EMBO J 35, 2566–2583, doi:10.15252/embj.201694253 (2016).

32 Pryde, K. R., Taanman, J. W. & Schapira, A. H. A LON-ClpP Proteolytic Axis Degrades Complex I to Extinguish ROS Production in Depolarized Mitochondria. Cell Rep 17, 2522–2531, doi:10.1016/j.celrep.2016.11.027 (2016).

33 Brodie, E. J., Zhan, H., Saiyed, T., Truscott, K. N. & Dougan, D. A. Perrault syndrome type 3 caused by diverse molecular defects in CLPP. Sci Rep 8, 12862, doi:10.1038/s41598-018-30311-1 (2018).

34 Jenkinson, E. M. et al. Perrault syndrome is caused by recessive mutations in CLPP, encoding a mitochondrial ATP-dependent chambered protease. Am J Hum Genet 92, 605–613, doi:10.1016/j.ajhg.2013.02.013 (2013).

35 Gispert, S. et al. Loss of mitochondrial peptidase Clpp leads to infertility, hearing loss plus growth retardation via accumulation of CLPX, mtDNA and inflammatory factors. Hum Mol Genet 22, 4871–4887, doi:10.1093/hmg/ddt338 (2013).

36 Ishizawa, J. et al. Mitochondrial ClpP-Mediated Proteolysis Induces Selective Cancer Cell Lethality. Cancer Cell 35, 721–737 e729, doi:10.1016/j.ccell.2019.03.014 (2019).

37 Wang, S. & Dougan, D. A. The Direct Molecular Target for Imipridone ONC201 Is Finally Established. Cancer Cell 35, 707–708, doi:10.1016/j.ccell.2019.04.010 (2019).

38 Nishimura, K. et al. Discovery of a Unique Clp Component, ClpF, in Chloroplasts: A Proposed Binary ClpF-ClpS1 Adaptor Complex Functions in Substrate Recognition and Delivery. Plant Cell 27, 2677–2691, doi:10.1105/tpc.15.00574 (2015).

39 Yeom, J. & Groisman, E. A. Activator of one protease transforms into inhibitor of another in response to nutritional signals. Genes Dev, doi:10.1101/gad.325241.119 (2019).

40 Dougan, D. A., Reid, B. G., Horwich, A. L. & Bukau, B. ClpS, a substrate modulator of the ClpAP machine. Mol Cell 9, 673–683 (2002).

41 Dougan, D. A., Weber-Ban, E. & Bukau, B. Targeted delivery of an ssrA-tagged substrate by the adaptor protein SspB to its cognate AAA+ protein ClpX. Mol Cell 12, 373–380 (2003).

42 Wojtyra, U. A., Thibault, G., Tuite, A. & Houry, W. A. The N-terminal zinc binding domain of ClpX is a dimerization domain that modulates the chaperone function. J Biol Chem 278, 48981–48990 (2003).

43 Truscott, K. N., Lowth, B. R., Strack, P. R. & Dougan, D. A. Diverse functions of mitochondrial AAA+ proteins: protein activation, disaggregation, and degradation. Biochem Cell Biol 88, 97–108, doi:10.1139/o09-167 (2010).

44 Mitchell, A. L. et al. InterPro in 2019: improving coverage, classification and access to protein sequence annotations. Nucleic Acids Res 47, D351–D360, doi:10.1093/nar/gky1100 (2019).

45 Watanabe, K., Yumimoto, K. & Nakayama, K. I. FBXO21 mediates the ubiquitylation and proteasomal degradation of EID1. Genes Cells 20, 667–674, doi:10.1111/gtc.12260 (2015).

46 Zhang, C. et al. Peptidic degron in EID1 is recognized by an SCF E3 ligase complex containing the orphan F-box protein FBXO21. Proc Natl Acad Sci U S A 112, 15372–15377, doi:10.1073/pnas.1522006112 (2015).

47 Montandon, C., Dougan, D. A. & van Wijk, K. J. N-degron specificity of chloroplast ClpS1 in plants. FEBS Lett 593, 962–970, doi:10.1002/1873-3468.13378 (2019).

48 Erbse, A. et al. ClpS is an essential component of the N-end rule pathway in Escherichia coli. Nature 439, 753–756, doi:10.1038/nature04412 (2006).

49 Puri, N. & Karzai, A. W. HspQ Functions as a Unique Specificity-Enhancing Factor for the AAA+ Lon Protease. Mol Cell 66, 672–683 e674, doi:10.1016/j.molcel.2017.05.016 (2017).

50 Varshavsky, A. N-degron and C-degron pathways of protein degradation. Proc Natl Acad Sci U S A 116, 358–366, doi:10.1073/pnas.1816596116 (2019).

51 Catanzariti, A. M., Soboleva, T. A., Jans, D. A., Board, P. G. & Baker, R. T. An efficient system for high-level expression and easy purification of authentic recombinant proteins. Protein Sci 13, 1331–1339, doi:10.1110/ps.04618904 (2004).

52 Kang, S. G. et al. Functional proteolytic complexes of the human mitochondrial ATP-dependent protease, hClpXP. J Biol Chem 277, 21095–21102, doi:10.1074/jbc.M201642200 (2002).

53 Krzysiak, T. C., Chen, B. B., Lear, T., Mallampalli, R. K. & Gronenborn, A. M. Crystal structure and interaction studies of the human FBxo3 ApaG domain. FEBS J 283, 2091–2101, doi:10.1111/febs.13721 (2016).

54 Shima, Y. et al. PML activates transcription by protecting HIPK2 and p300 from SCFFbx3-mediated degradation. Mol Cell Biol 28, 7126–7138, doi:10.1128/MCB.00897-08 (2008).

55 Mallampalli, R. K. et al. Targeting F box protein Fbxo3 to control cytokine-driven inflammation. J Immunol 191, 5247–5255, doi:10.4049/jimmunol.1300456 (2013).

56 Levchenko, I., Grant, R. A., Wah, D. A., Sauer, R. T. & Baker, T. A. Structure of a delivery protein for an AAA+ protease in complex with a peptide degradation tag. Mol Cell 12, 365–372 (2003).

57 Levchenko, I., Seidel, M., Sauer, R. T. & Baker, T. A. A specificity-enhancing factor for the ClpXP degradation machine. Science 289, 2354–2356 (2000).

58 Park, E. Y. et al. Structural basis of SspB-tail recognition by the zinc binding domain of ClpX. J Mol Biol 367, 514–526, doi:10.1016/j.jmb.2007.01.003 (2007).

59 Truscott, K. N. et al. A presequence- and voltage-sensitive channel of the mitochondrial preprotein translocase formed by Tim23. Nat Struct Biol 8, 1074–1082 (2001).

60 Zheng, L., Baumann, U. & Reymond, J. L. An efficient one-step site-directed and site-saturation mutagenesis protocol. Nucleic Acids Res 32, e115, doi:10.1093/nar/gnh110 (2004).

61 Ninnis, R. L., Spall, S. K., Talbo, G. H., Truscott, K. N. & Dougan, D. A. Modification of PATase by L/F-transferase generates a ClpS-dependent N-end rule substrate in Escherichia coli. EMBO J 28, 1732–1744, doi:10.1038/emboj.2009.134 (2009).

62 Schagger, H. & von Jagow, G. Tricine-sodium dodecyl sulfate-polyacrylamide gel electrophoresis for the separation of proteins in the range from 1 to 100 kDa. Anal Biochem 166, 368–379 (1987).

63 Geissler, A. et al. The mitochondrial presequence translocase: an essential role of Tim50 in directing preproteins to the import channel. Cell 111, 507–518 (2002).

64 Bezawork-Geleta, A., Saiyed, T., Dougan, D. A. & Truscott, K. N. Mitochondrial matrix proteostasis is linked to hereditary paraganglioma: LON-mediated turnover of the human flavinylation factor SDH5 is regulated by its interaction with SDHA. FASEB J 28, 1794–1804, doi:10.1096/fj.13-242420 (2014).

65 Stojanovski, D., Pfanner, N. & Wiedemann, N. Import of proteins into mitochondria. Methods Cell Biol 80, 783–806, doi:10.1016/S0091-679X(06)80036-1 (2007).

66 Kabsch, W. Integration, scaling, space-group assignment and post-refinement. Acta Crystallographica Section D 66, 133–144, doi:doi:10.1107/S0907444909047374 (2010).

67 Kabsch, W. XDS. Acta Crystallographica Section D 66, 125–132, doi:doi:10.1107/S0907444909047337 (2010).

68 Afonine, P. V. et al. Joint X-ray and neutron refinement with phenix.refine. Acta Crystallogr D Biol Crystallogr 66, 1153–1163, doi:10.1107/S0907444910026582 (2010).

