## Supplementary for "PDIP38 is a novel adaptor-like modulator of the mitochondrial AAA+ protease CLPXP"

**Supplementary Table 1 Data collection and refinement statistics**

| | PDIP38 |
|---|---|
| **Data collection** | |
| Space group | $P6_2$ |
| Cell dimensions | |
| *a*, *b*, *c* (Å) | 120.1, 120.1, 49.2 |
| α, β, γ (°) | 90, 90, 120 |
| Resolution (Å) | 50 – 3.08 (3.26 – 3.08) |
| $R_{sym}$ or $R_{merge}$ | 0.07 (2.62) |
| CC* in outermost shell | 12.2 |
| $I / \sigma I$ | 11.2 (0.59) |
| Completeness (%) | 97.0 (94.6) |
| Redundancy | 3.95 (3.95) |
| **Refinement** | |
| Program | PHENIX |
| Resolution (Å) | 50 – 3.08 (3.34 – 3.08) |
| No. reflections | 6006 |
| $R_{work}$ / $R_{free}$ | 0.26/0.28 (0.31/0.34) |
| No. atoms | |
| Protein | 2087 |
| *B*-factors | |
| Protein | 28.2 |
| R.m.s. deviations | |
| Bond lengths (Å) | 0.003 |
| Bond angles (°) | 0.67 |
| **Ramachandran statistics** | |
| Residues in favored region No (%) | 91 |
| Residues in allowed region No (%) | 8.5 |
| Residues in outlier region No (%) | 0.4 |
| **PDB-entry** | 6SNM |

*Values in parentheses are for highest-resolution shell.

**Supplementary Table 2 Conserved residues that line and cap the Hydrophobic groove**

| Human PDIP38 | Human Fbx3 (5HDW) | *X. axonopodis* ApaG (2F1E) |
|---|---|---|
| Y 265 | Y 308 | Y 33 |
| R 282 | R 329 | R 50 |
| W 284 | W 331 | W 52 |
| I 286 | I 333 | I 54 |
| E 294 | E 341 | E 62 |
| V 296 | V 343 | V 64 |
| V 301 | V 348 | V 69 |
| V 302 | V 349 | V 70 |
| Y 317 | Y 363 | Y 84 |
| V 321 | T 367 | V 88 |
| M 330 | M 376 | M 97 |

**Supplementary Table 3 Oligonucleotide primers used in this study**

| Primer | DNA sequence (5'-> 3') | Gene | Features |
|---|---|---|---|
| 5pdip_sac2 | GACTCTCCGCGGTGGATCCTCC CGAAACCGACCAGAGGGC | PDIP38 | Sac II restriction site for cloning into pHUE |
| 3pdip_hind | GCTACGAAGCTTCTACCAGTGA AGGCCTGAGGGTGG | PDIP38 | Hind III restriction site for cloning into pHUE |
| 5pdip_not | GCAGTAGCGGCCGCATCGTCCC GAAACCGACCAGAG | PDIP38 | Not I restriction site for cloning into pET10N |
| 5not_Cdip | GCAGTAGCGGCCGCACGGGAAA CAACTGAGAACATACG | PDIP38$_C$ | Not I restriction site for cloning into pET10N |
| 3pdip_xho | GATAGCCTCGAGCTACCAGTGA AGGCCTGAGGGTGG | PDIP38 PDIP38$_C$ | Xho I restriction site for cloning into pET10N or pGEX4T |
| 5pdip_bam | GTCGATGGATCCTCCCGAAACC GACCAGAGGGC | PDIP38 | Bam HI restriction site for cloning into pGEX4T |
| Ndip_3not | ACGTAGCGCGGCCGCATGAACA TCGGAGAGCTCCAG | PDIP38$_N$ | Not I restriction site for cloning into pET10C |
| 5pdip_nde | CGTATCCATATGTCCTCCCGAA ACCGACCAGAGGGC | PDIP38 PDIP38$_N$ | Nde I restriction site for cloning into pET10C |
| 3pdip_not | GATAGCTGCGGCCGCCCAGTGA AGGCCTGAGGGTGG | PDIP38 PDIP38$_C$ | Not I restriction site for cloning into pET10C |
| 3dip_STOP | GATAGCTGCGGCCGCCTACCAG TGAAGGCCTGAGGGTGG | PDIP38 | Not I restriction site for cloning into pET10C (no His tag) |
| Ndip_3xho | ACGTAGCTCGAGCTAATGAACA TCGGAGAGCTCCAG | PDIP38$_N$ | Xho I restriction site for cloning into pGEX4T |
| PDIP_bam1 | CTCAGACAGAATAAGGATCCTT CTTGGCTAACCATG | PDIP38$_N$ PDIP38$_C$ | Introduce stop codon and Bam HI restriction site to create PDIP38$_N$ or PDIP38$_C$ in pGEX4T |
| PDIP_bam2 | GCCAAGAAGGATCCTTATTCTG TCTGAGATCTCTGAG | PDIP38$_N$ PDIP38$_C$ | Introduce stop codon and Bam HI restriction site to create PDIP38$_N$ or PDIP38$_C$ in pGEX4T |
| 5prePDIP_hind3 | TCGTAGAAGCTTATGGCAGCCT GTACAGCCCGGC | PDIP38 | Hind III restriction site for cloning into pEGFP-N1 |
| 38G2 | CGGTGGATCCCAGTGAAGGCCT GAGGGTG | PDIP38 | Bam HI restriction site for cloning into pEGFP-N1 |
| hX4A_1 | CTTTGTAGCTGCAGCCGCTTTT GTCAAGTGTGAAAAG | CLPX CLPX$_N$ | Introduce Pst I restriction site (when converting wild type CLPX to CLPX$_{AAAA}$ |
| hX4A_2 | CTTGACAAAAGCGGCTGCAGCT ACAAAGGTCTCTAC | CLPX CLPX$_N$ | Introduce Pst I restriction site (when converting wild type CLPX to CLPX$_{AAAA}$ |
| Sac2_TRALP | CGAATTCCGCGGTGGAACCCGG GCTCTCCCGCTCATTC | CLPP | Sac II restriction site for cloning into pHUE |
| LhP_hind3 | GCTACGAAGCTTAGGTGCTAGC TGGGACAGGTTC | CLPP | Hind III restriction site for cloning into pHUE |

[1] restriction sites for cloning and/or screening are underlined

**Supplementary Table 4 Plasmids used in this study**

| Plasmid name | Plasmid description | Plasmid features, source |
|---|---|---|
| pDT1329 | pOTB7/*PDIP38* | I.M.A.G.E. clone 3349399 |
| pDT1432 | pHUE/PDIP38 | Amplified *PDIP38* using 5pdip_sac2 and 3pdip_hind, digested with *Bam* HI and *Hind* III and cloned into pHUE |
| pDT1355 | pET10N/*PDIP38* | Amplified *PDIP38* using 5pdip_not and 3pdip_xho, digested with *Not* I and *Xho* I and cloned into pET10N |
| pDT1562 | pET10C/*PDIP38* | Amplified *PDIP38* using 5pdip_nde and 3pdip_not, digested with *Nde* I and *Not* I and cloned into pET10C |
| pDT1586 | pET10C/*PDIP38* | Amplified *PDIP38* using 5pdip_nde and 3dip_STOP, digested with *Nde* I and *Not* I and cloned into pET10C |
| pDT1356 | pGEX-4T/*PDIP38* | Amplified *PDIP38* using 5pdip_bam and 3pdip_xho, digested with *Bam* HI and *Xho* I and cloned into pGEX-4T |
| pDT1367 | pGEX-4T/PDIP38$_N$ | Quick change mutagenesis using pDT1356 and primers PDIP_bam1 and PDIP_bam2 |
| pDT1362 | pGEX-4T/PDIP38$_C$ | Quick change mutagenesis using pDT1356 and primers PDIP_bam1 and PDIP_bam2, digestion with *Bam* HI to remove the fragment coding for *PDIP38*$_N$ followed by ligation of digested plasmid |
| pDT2191 (-94) | pE-FLAG/PDIP38-FLAG | Amplified *PDIP38* using 5prePDIP_hind3 and 38G2, digested with *Hind* III and *Bam* HI and cloned into pE-FLAG |
| pDT1766 (-67) | pNHIS/$H_6$GFP-PDIP38 | Subcloned from pDT1355 (digestion with *Not* I and *Hind* III) |
| pDT1279 | pET10C/*CLPX* | Lowth et al., 2012 |
| pDT1255 | pET10C/*CLPX*$_{ZBD}$ | Lowth et al., 2012 |
| pDT1260 | pET10C/*CLPX*$_E$ | Lowth et al., 2012 |
| pDT1411 | pET10C/*CLPX*$_{WB}$ | Lowth et al., 2012 |
| pDT1973 | pET10C/*CLPX*$_{4A}$ | Quick change mutagenesis using pDT1279 and primers hX4A_1 and hX4A_2 |
| pDT1977 | pET10C/*ZBD*$_{4A}$ | Quick change mutagenesis using pDT1255 and primers hX4A_1 and hX4A_2 |
| pDT1668 | pUHIS/*CLPP* | Lowth et al., 2012 |
| pDT2772 | pHUE/*CLPP* | Bezawork-Geleta et al., 2013 |
| pDD795 | pC10HIS/ec clpX$_{ZBD}$ | Dougan et al., 2003 |

### Supplementary Figure Legends

***Supplementary Figure 1. The in vitro degradation of FITC-$\alpha_{S2}$-casein by CLPXP is inhibited by PDIP38.***

The rate of FITC-$\alpha_{S2}$-casein degradation (white bars) was determined in the absence (column 1) or presence of increasing concentrations of PDIP38 [1.2 μM (column 2), 4.8 μM (column 3), 9.6 μM (column 4), 19.2 μM (column 5), 38.4 μM (column 6)]. The rate of FITC-κ-casein degradation (black bars) was determined in the absence (column 7) or presence of 38.4 μM PDIP38 (column 8).

***Supplementary Figure 2. The steady state levels of CLPX are reduced in cells lacking PDIP38.***

**a**. Full length slab of Figure 1b illustrating the specificity of siRNA mediated knock down and PDIP38 antisera. Proteins were separated by 15% Tris-glycine SDS-PAGE and subjected to immunoblotting with the appropriate antisera to visualize endogenous proteins. (*, non-specific cross-reactive protein in PDIP38 antisera, upper band on 15% Tris-glycine SDS-PAGE). **b.** The steady state levels of PIDP38 (top panel), CLPX ($2^{nd}$ panel) SDHA ($3^{rd}$ panel) and GAPDH (bottom panel) were analysed in HeLa cells 72 hours post-transfection with either Silencer Select Negative Control No. 1 siRNA (nc1, lane 1) and Negative Control No. 2 (nc2, lane 2) and compared to HeLa cells treated with *PDIP38*-targeted Silencer Select siRNAs s25055 (s55, lane 3) and s25056 (s56, lane 4). **c.** The steady state levels of PIDP38 were analysed in HeLa cells 72 hours post-transfection with either Silencer Select Negative Control No. 1 siRNA (nc1, lane 2) and Negative Control No. 2 (nc2, lane 4) and compared to HeLa cells treated with *PDIP38*-targeted Silencer Select siRNAs s25055 (s55, lane 3) and s25056 (s56, lane 5) and compared to PIDP38 levels in 20 μg (lane 6) or 40 μg (lane 7) mitochondria (mito). (**b-c**) Proteins were separated by 16.5% Tris-Tricine SDS-PAGE and subjected to immunoblotting with the appropriate antisera to visualize endogenous proteins. (*, non-specific cross-reactive protein in PDIP38 antisera, lower band on 16.5% Tris-Tricine SDS-PAGE).

***Supplementary Figure 3. PDIP38 does not inhibit the LONM-mediated degradation of casein***

*In vitro* degradation of casein by $LONM_6$ protease (400 nM) in the absence (lane 2 – 7) or presence of 1 μM PDIP38 (lanes 8 – 13). Proteins were separated by 10% Tris-Tricine SDS-PAGE and visualised by CBB staining.

***Supplementary Figure 4. Structure of PDIP38 yccV-like domain, relative to DUF525 domain***

Ribbon representation of human PDIP38 N-terminal yccV-like domain highlighting the β-strands (purple) and loops (tans), with C-terminal domain shown in surface representation. Loop 3 (L3) and loop 4 (L4) are disordered and hence represented by a dotted line. The extended β2 and β3 strands form a continuous sheet with the tope sheet of the Immunoglobulin-like fold of the DUF525 domain. The figure was generated in ChimeraX_Daily.

***Supplementary Figure 5. Structural alignment of PDIP38 with Fbx3 and ApaG highlighting the conserved hydrophobic residues of the groove.***

**a.** Ribbon representation of human PDIP38 (blue), aligned with the DUF525 domain of Fbx3 (green) and ApaG (red) illustrating the conserved hydrophobic residues that line the putative binding groove. **b.** Protein sequence alignment of human PDIP38 illustrating the absolutely conserved residues (bold) that are located within the hydrophobic pocket residues (red). **c.** Ribbon representation of DUF525 domain of Fbx3 (green). **d**. Ribbon representation of DUF525 domain of ApaG (red).

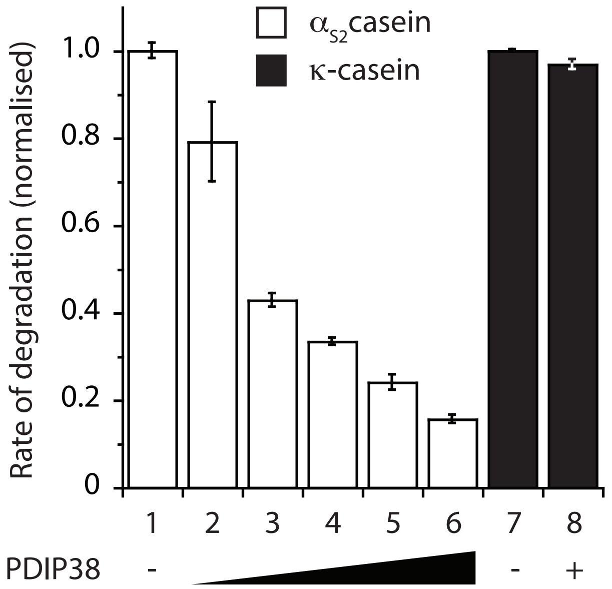


Strack et al., Supplementary Figure 1

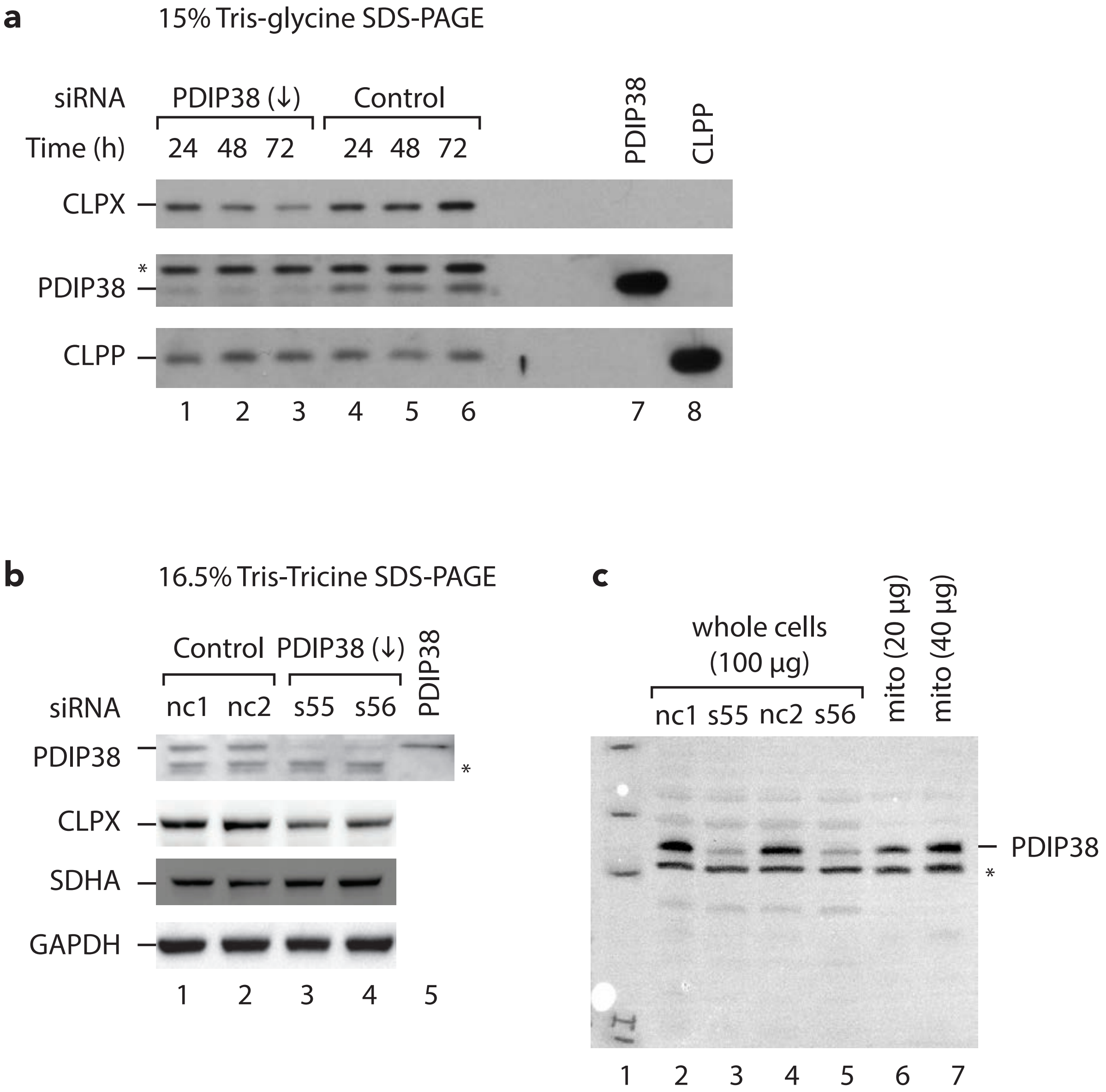


Strack et al., Supplementary Figure 2

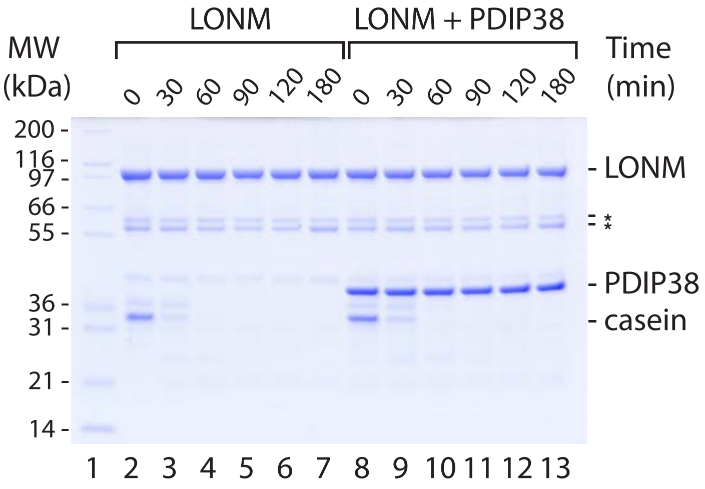


Strack et al., Supplementary Figure 3

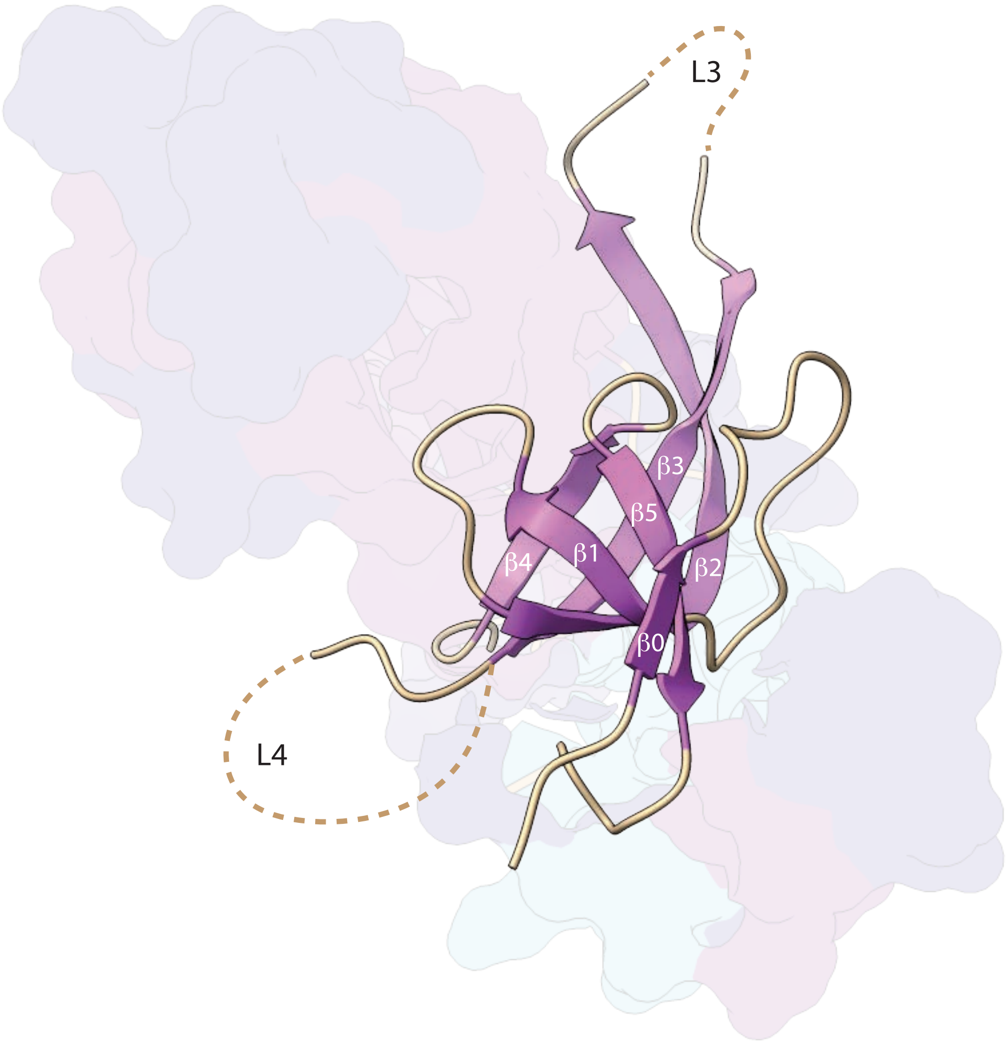


Strack et al., Supplementary Figure 4

**a**

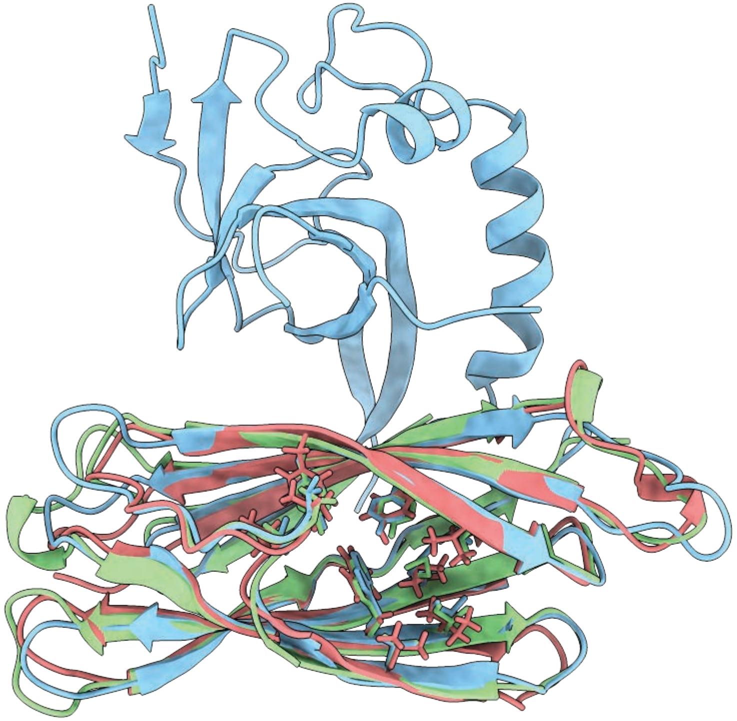

**b**

```
PDIP38      DVHRETTENIRVTVIPFYMGMREAQNSHVYWWRYC
Fbx3        SEFVATTGDITVSVSTSFLPELSSVHPPHYFFTYR
ApaG        ------RYRVEVEVSPRFLAHQSTPDEGRYAFAYS
Consensus   ------p-plpVpV-s-ahs-ps--p---Yha-Y-

PDIP38      IRLENLDS----DVVQLRERHWRIFSLSGTLETVR
Fbx3        IRIEMSKDALPEKACQLDSRYWRITNAKGDVEEVQ
ApaG        IRIQNAGA----VPARLVARHWQITDGNGRTEQVD
Consensus   IRlp---s-----shpL--RhWpIhs-pGphEpVp

PDIP38      GRGVVGREPVLSKEQPAFQYSSHVSLQASSGHMWG
Fbx3        GPGVVGEFPIISPGR-VYEYTSCTTFSTTSGYMEG
ApaG        GEGVVGEQPWLRPGE-AFHYTSGVLLETEQGQMQG
Consensus   G-GVVGcbPhlp--p-hapYoS-h-hphppG-MbG

PDIP38      TFRFERPD--GSHFDVRIPPFSLESNKDEKTPPSG
Fbx3        YYTFHFLYFKDKIFNVAIPRFHMACPT--------
ApaG        HYDMVAD--DGTEFIAPIAAFVLS-----------
Consensus   haph------sp-F-h-Is-F-h------------
```

**c**

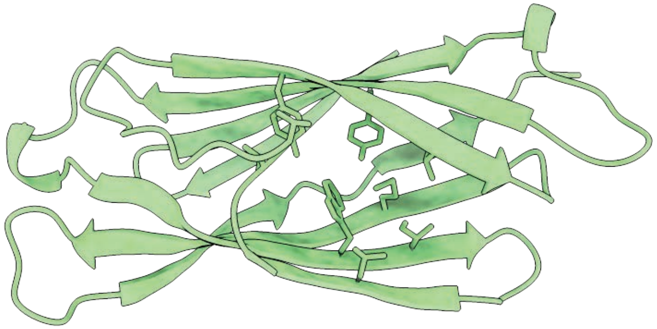

Human Fbx3 DUF525 domain (5HDW)

**d**

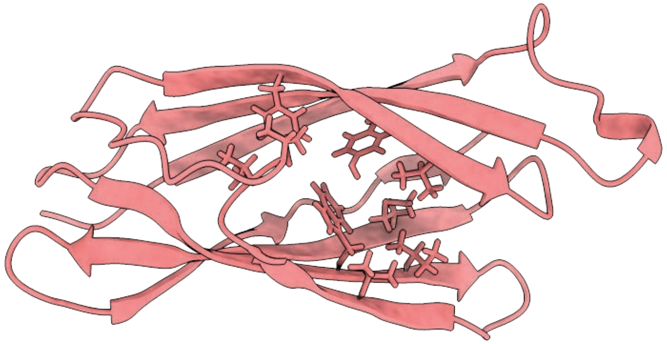

Xanthomonas axonopodis ApaG (2F1E)

Strack et al., Supplementary Fig 5
